# Experience reorganizes content-specific memory traces in macaques

**DOI:** 10.1101/2025.04.08.647787

**Authors:** Saman Abbaspoor, Ayman Aljishi, Kari L. Hoffman

**Author notes:** Present address: F.M. Kirby Neurobiology Center and Rosamund Stone Zander Translational Neuroscience Center, Boston Children’s Hospital; Department of Neurology, Harvard Medical School; Boston, MA, USA.

## Abstract

Memory formation relies on the reorganization of neural activity patterns during experience that persist in subsequent sleep. How these processes promote learning while preserving established memories remains unclear. We recorded neural ensemble activity from hippocampal and associated regions in freely moving macaques as they recalled item sequences presented that day (“new”), one day prior (“recent”), or over two weeks prior (“old”). Cell assemblies biased for old sequences showed less drift, greater network connectivity, and stronger sleep reactivation than new-biased assemblies. Pairs of old and recent assemblies formed persistent task-to-sleep coupling (“metassemblies”), unlike new assembly pairs. In the hippocampus, the propensity for superficial and deep CA1 pyramidal cells to form integrated assemblies increased with memory age. These findings reveal rapid organization and stabilization of neural activity in the primate brain, suggesting potential mechanisms for balancing learning with memory linking and durability.

## Main

Memory formation and organization are dynamic, ongoing processes that allow the brain to encode, update, and integrate multiple experiences over time. In rodents, accumulating evidence shows that recently formed memories are initially supported by selective activation of neuronal ensembles during learning and subsequently stabilized through their reactivation during offline states such as sleep^1–6^. Similar reactivation patterns have been observed in other species including birds^7^, bats^8^, non-human primates^9,10^ and, more recently, in humans^11–13^. Although, these findings suggest that some of the hippocampal circuit mechanisms are broadly similar across species, differences in ethological demands and behavioral repertoire may shape both the content of reactivated patterns—what types of experiences are prioritized for reactivation and retention— and the organization of temporally distinct experiences into an adaptive memory system. Given that much of the existing work has focused on the maintenance of stable representations of discreet, isolated, and/or recent memories, it remains unclear how the brain manages to keep track of multiple distinct experiences over time and organizes memories originating from different points in time^14–18^. This question is particularly relevant in primates, whose extended lifespans and cognitive flexibility require memory systems that support not only the preservation of individual experiences but also their integration into coherent, behaviorally adaptive representations. Whereas waking experience may support and recruit some of these processes^19–25^, offline states and specifically sleep are thought to play a crucial role in this reorganization^26,27^. In this study, we sought to address whether consistent activity patterns reflecting multiple spatiotemporal events (memoranda) are reactivated during sleep in macaques, and if so, how the patterns of activation and reactivation may vary as a function of prior experience. Using high-density wireless recordings from hippocampal and extrahippocampal regions, we tracked experience-dependent changes in ensemble activity that could hold key insights into how the primate brain preserves older memories while incorporating new ones.

## Newly-learned and remotely-learned sequences can be decoded from synchronized spiking ensembles

We used high-density linear arrays to simultaneously record the single unit activity from neuronal ensembles in the hippocampus and connected extrahippocampal regions of two freely-moving macaques (range: 31-273 units per session, N = 35 sessions, Fig. 1c-e and Extended Data Fig. 1b-d). Task recordings took place in the ‘*Treehouse*’—a touchscreen-enabled 3D enclosure, followed by sleep recordings taken overnight in their housing room (Fig. 1a and Extended Data Fig. 1a). ‘Day 1’ Treehouse sessions were comprised of two types of repeated ‘Z’ shaped trajectories: a ‘New’, 4-item-context sequence spanning the 4 touchscreens of one corner, and an ‘Old’ sequence presented in the opposite corner that had been learned in that corner, on average, 6 weeks prior with no intervening exposure (Fig. 1a and Extended Data Fig. 1b-c, see Methods). This allowed simultaneous recording during performance of two sequences that differed in their prior exposure, but with the corner location of the Old/New sequence types counterbalanced across sets (Fig. 1c-e, Extended Data Fig. 2). Individual learning results in this task have been reported previously^28^. Accuracy was higher for Old sets than New, indicating long-term memory savings (Fig. 1b, p < 0.001, permutation test with FDR correction). ‘Day 2’ sessions occurred the next day and used the same stimulus sets and assigned locations as Day 1, but with the New set now termed ‘Recent’. Day 2 performance on Recent sets improved (p < 0.001, permutation test with FDR correction), while remaining below Old set levels, suggesting overnight retention of the newly-learned sequences and a residual benefit of long-term, remote memory for the old sequences. To determine whether the recorded neural activity could contribute to selective memory for these sequences, we trained a linear support vector machine (SVM) decoder to classify trial identity (Old vs. New/Recent) based on single-unit or ensemble activity (Fig 1g and Extended Data Fig. 3). Decoding based on ensemble activity consistently outperformed decoding that used single unit activity, and it surpassed shuffling controls (Fig. 1g, Extended Data Fig. 3). Movement-related variables during the task, including angular velocity and linear acceleration, did not differ significantly between the new and old sequences performed in their respective corners and are therefore unlikely to account for the observed decoding performance (Extended Data Fig. 1d-e).These results indicate that the pattern of spiking activity in the recorded ensemble generally varied as a function of trial identity.

**Fig 1.**
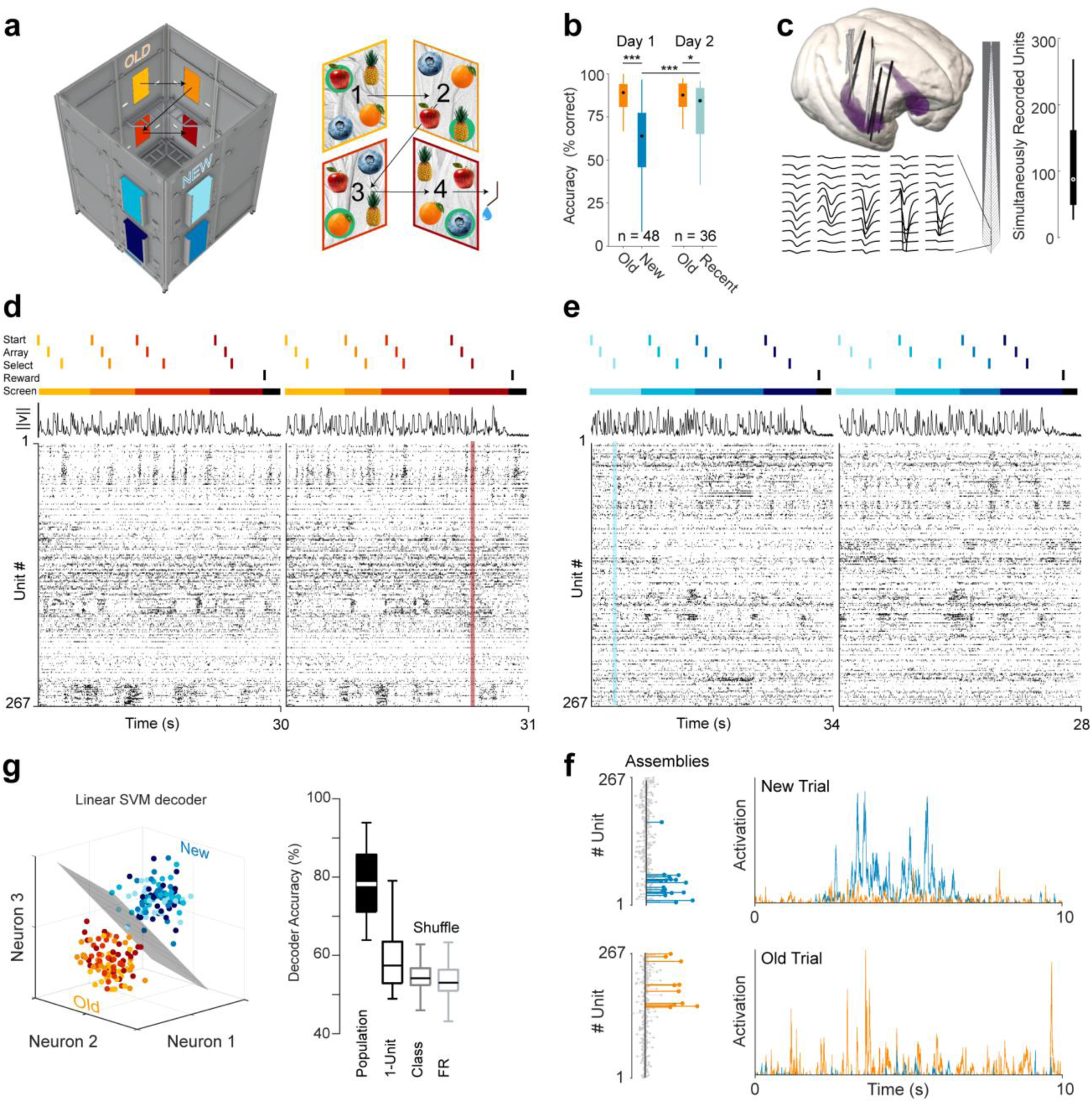
Experimental design and neural population analysis methods **(a)** (*Left*) Three-dimensional rendering of the Treehouse 3-D task environment indicating the opposite corners where “old” and “new” visual stimulus arrays are presented sequentially on touchscreens. (*Right*) Trial schematic illustrating an example item-context sequence across four screens, with the correct target items circled in green. Item positions on the screens are randomized across screens and trials. Old and new item-context sequences (corners) are visited in alternating blocks (see Methods and Ref^28^) **(b)** Target selection rates per sequence condition. Performance of new sequences improved between the first and second days of presentation (permutation test with FDR correction, p < 0.001), and performance of old sequences exceeded newly-presented sequences (vs. ‘New’ first day, permutation test with FDR correction p < 0.001, vs. ‘Recent’ Second day permutation test with FDR correction p < 0.05), trial category label permutation tests.) **(c)** (*Left*) CT-MR reconstructed electrode trajectories for Monkey 1 (black) and Monkey 2 (gray) projected to NIMH macaque template brain^34^, with representative spike waveforms shown across adjacent recording channels from the high-density arrays. (*Right*) Isolated single unit yields across sessions (Monkey 1: n=17 sessions, Monkey 2: n=18 sessions). **(d)** *(Top)* Timing of trial events across a sequence of four screens for two “old” trials. *(Middle)* Movement trace, shown as the vector norm of angular velocity. *(Bottom)* Raster plots of corresponding population spiking activity (time x cells), sorted according to the Rastermap method ^35^; an example ‘population vector’ of activity highlighted in red. **(e)** As in (D), but for two “new” trials from the same session. **(f)** *(Left)* Schematic of linear SVM decoding of trial type (Old/New) based on population vector activity. (*Right*) Decoder accuracy of trial epoch identity classification for different SVM models (n=26 population; 3332 single units)**. (g)** Task-selective cell assemblies (*Left*) Examples of cell assemblies from the same session, with member weights colored according to biased activation for old (orange) or new (blue) conditions. (*Right*) Activation strengths of both assemblies (blue, orange, respectively) presented for a new-sequence trial (top) and an old-sequence trial (bottom) demonstrating biased activation by trial.

We used component analysis (PCA/ICA, see methods^29–31^) to identify distinct ensembles of coactive neurons (Fig. 1f, Extended Data Fig. 4), as an operational definition of “cell assemblies” (see also Fig 1f and Extended Data Fig. 4A). We fit a generalized linear model (GLM) to each detected assembly to assess how well activation strength could be predicted by specific trial elements, such as sequence position, behavioral outcomes (e.g., trial success), and movement parameters. This approach also allowed us to examine which factors tended to covary across assemblies (Extended Data Fig 5 and 6). To determine whether these cell assemblies reappear during post-learning sleep and whether offline processing differed between assemblies biased toward new versus old information, we grouped cell assemblies into Old, New/Recent, or Nonselective categories based on their activation likelihood between the two stimulus sequences. Notably, these categories of assemblies did not differ in quality metrics (OtsuEM^32,33^) or in the firing rates of their member neurons, indicating that key features relevant for detecting and defining cell assemblies were comparable across groups. Old– and Recent-biased assemblies were, however, moderately larger (i.e. had more members) than New-biased assemblies (Extended Data Fig. 4b).

## Experience biases sleep reactivation of task-related cell assemblies

We tracked assembly activation during task and reactivation in sleep and found that all categories of cell assemblies reactivated in sleep (Fig. 2a and Extended Data Fig. 7). Old– and Recent-biased assemblies showed greater and more frequent activation and reactivation than new-biased assemblies (Fig. 2c, and Extended Data Fig. 7a, p < 0.001, permutation test with FDR correction). This gain in the reactivation strength was not due to changes in the average firing rate of the members (Fig. 2e). All three task groups showed linear correlation between task activation and sleep ripple reactivation (Fig 2b); however, this correlation was stronger for Old and Recent cell assemblies compared to New assemblies, with no difference between Old and Recent (Fig. 2b, p < 0.01 Spearman correlation with permutation test, and Fisher’s z test with FDR correction, p < 0.001). In addition to the preservation of relative assembly strength into sleep, both New and Old assemblies showed a reactivation recency bias: they were more strongly reactivated when their respective sequence was the last one experienced before sleep (Fig 2d). This recapitulates findings from rodent studies ^22,36,37^ and reinforces the idea that both new and old assemblies are well formed, yet their reactivation in sleep is subject to task-related influence.

**Fig 2.**
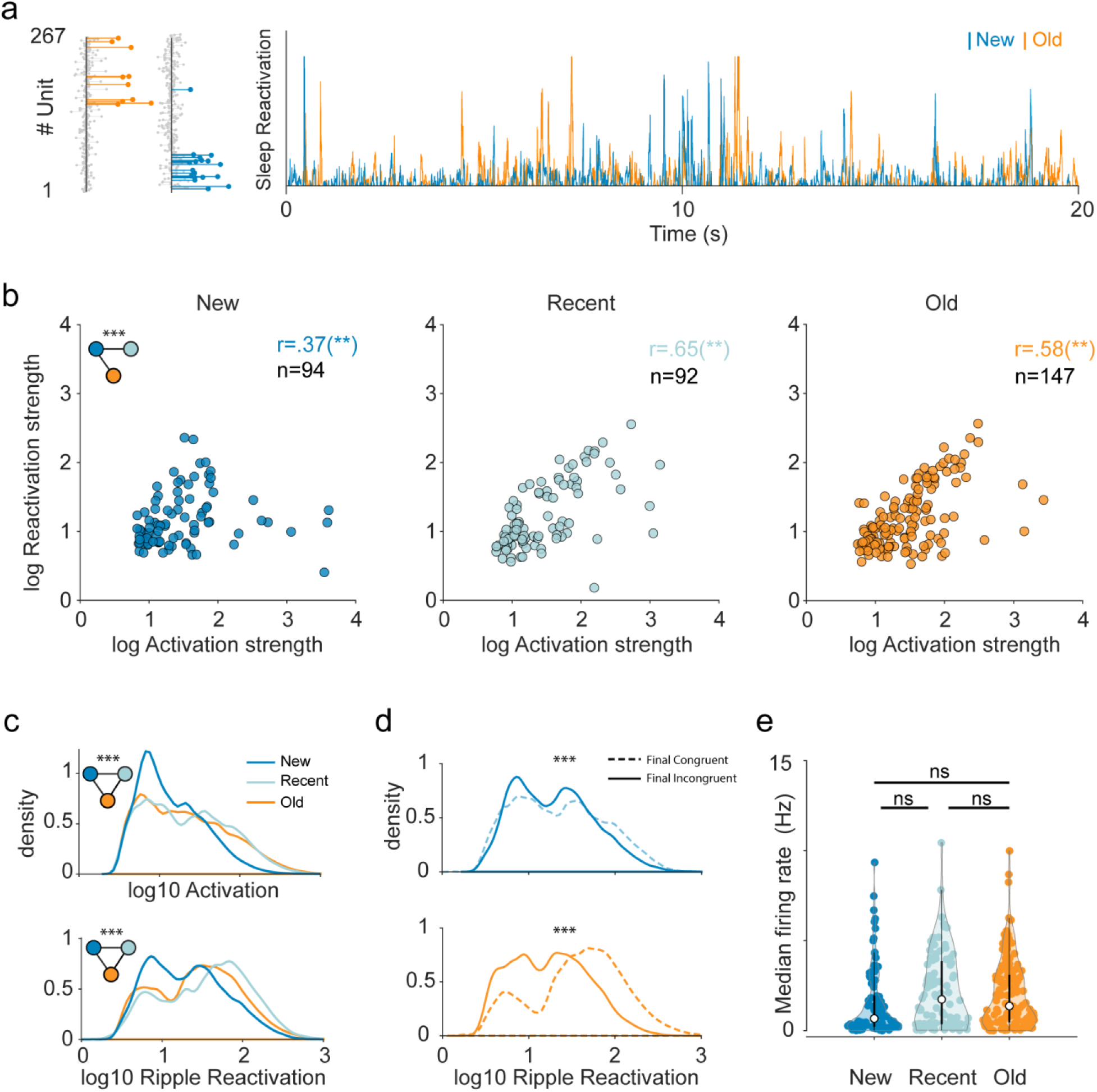
Selective reactivation of cell assemblies during sleep **(a)** Sleep reactivation strengths of two example cell assemblies obtained from task activation (Fig. 1F). **(b)** Scatterplots of mean sleep ripple reactivation strength per assembly as a function of mean task activation strength. Left plot shows new-sequence biased assemblies, middle plot the recent-sequence assemblies, and the right plot the old-sequence assemblies. Inset, left: Black line connecting dots indicate significant differences in activation-reactivation correlation between cell assembly categories as represented by dot color (p < 0.01 Spearman correlation with permutation and p < 0.001 Fisher’s z test with FDR correction). **(c)** Distributions of task activation (top) and ripple reactivation (bottom) strengths pooled across assemblies of the respective categories (new = blue, recent = light blue, orange = old). Inset: *** p < 0.001 pairwise distribution differences indicated by black lines connecting color-coded circles, permutation test). **(d)** Distributions of reactivation strength for New (top) and Old (bottom) assemblies, shown separately based on the Old/New context of the last trial block. ‘Final Congruent’ refers to cases where the context of the last trial block matches the activation bias of the cell assembly. ‘’Final Incongruent’ refers to cases where the final context and activation bias are mismatched (*** p < 0.001, pairwise permutation test). **(e)** Median firing rate across members of a cell assembly, grouped by assembly category. Permutation test with FDR correction.

## Assemblies link and stabilize over time

Emerging evidence indicates that, at the neuronal level, the hippocampus may support flexible, higher-order linking of smaller neuronal assemblies into larger integrated networks. Such linking could facilitate rapid learning and the efficient encoding of complex experiences^38^. Offline processes during rest or sleep may play a critical role in enabling such associations. If a pair of cell assemblies are linked, the correlation between their activation strength vectors should increase; if this coupling is stable, the correlation should remain consistent across time and behavioral states. To assess this possibility, we computed the correlation between pairs of assembly activations (i.e. Assembly coupling, Fig 3a). The correlations between New-biased assembly pairs, as well as between New– and Old-biased assemblies, increased during sleep, while correlations between other assembly types remained unchanged (Fig 3a). Importantly, when we sorted assembly coupling during task and sleep ripple epochs by task epoch, pairs of assemblies with recent and old memory biases exhibited greater stability—evident by high correlations between task and sleep—whereas coupling among New–New and New–Old pairs showed greater reorganization (Fig 3a). We observed the same pattern of results even after excluding assembly pairs with high cosine similarity (Extended Data Fig. 8d), confirming that the observed changes in coupling were not driven by redundant assembly membership (i.e. composition), but rather reflect genuine experience-dependent reorganization of inter-assembly dynamics. Changes in higher-order cell assembly coupling could stem from the cofiring changes of constituent neurons. We calculated cofiring as the pairwise correlation between neuron firing rates, comparing task vs. sleep cofiring within ripples to assess stability across states (Fig 3b and Extended Data Fig. 8 and 9). Average cofiring was greater for members of Old and Recent assemblies compared to New assemblies (p < 0.001, permutation test with FDR correction, Extended Data Fig. 9). These results suggest that sleep facilitates the emergence of higher-order interactions among assemblies, which stabilize with experience. The formation of such metassemblies—hierarchically organized groups of assemblies with varying degrees of association—may support flexible reorganization of assemblies for encoding new memories, while preserving core “elemental” assemblies linked to older experiences. Selective linking of assembly subsets could further enable the integration of new memories into pre-existing mnemonic frameworks.

**Fig 3.**
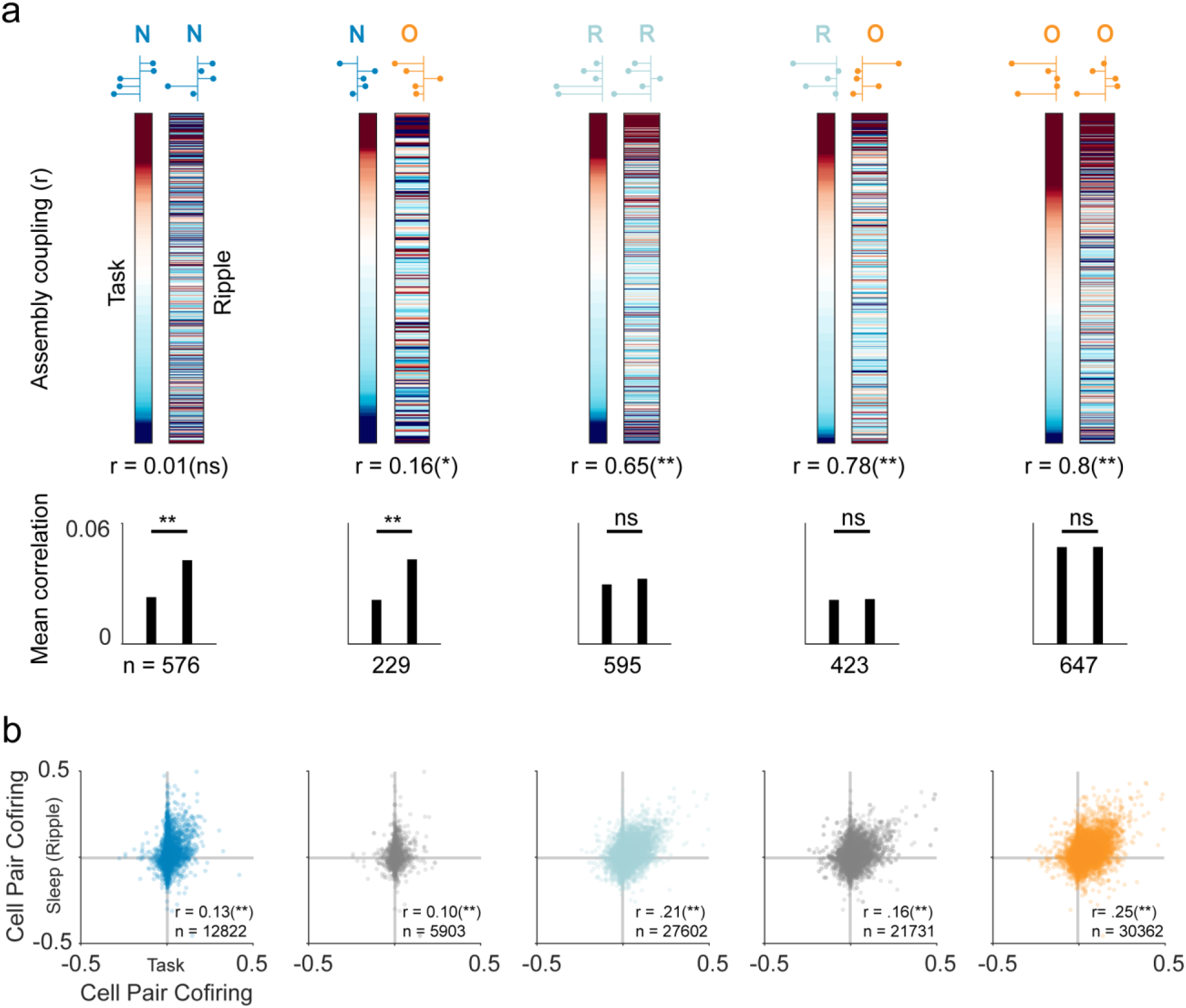
Changes in assembly coupling with memory age **(a)** (Top) Color-coded plots of coupling strength (co-re/activation) between pairs of cell assemblies across different memory categories during task events and sleep ripple events. Coupling was quantified as the Pearson’s correlation between pairs of re/activation traces. Pairs are sorted based on task activation coupling strength. The consistency of coupling across task and ripple epochs is quantified as the Pearson’s correlation shown below each plot. (Bottom) Mean coupling strength during task and sleep ripple events (* = p < 0.05, ** = p < 0.01, label permutation test with 10000 permutations). N= total number of assembly pairs per group. **(b)** Scatterplot of cofiring of pairs of neurons during sleep ripple events as a function of cofiring during task. Cell-pair cofiring was calculated as Spearman’s rank correlation between spike trains for each neuron pair. The consistency of coupling between task and sleep co-firing is quantified with Pearson’s correlation (r).

## Network mechanisms of cell assembly dynamics

Changes from task activation to sleep reactivation, cell assembly coupling, and pairwise neuronal cofiring indicate that cell assemblies undergo reorganization based on experience. Yet most methods for assessing assembly reactivation or replay assume a stationary activation “template” at least within the reference session^5,39^. In particular, the PCA/ICA approach defines assemblies statically and assumes temporal stationarity within the reference session, failing to capture changes in the structure of neuronal correlations over time. Recent findings suggest that neural ensembles dynamically reorganize to update memories and track ongoing experience^40–45^. In our study, if cell assemblies reorganize through changes in the firing rates of participating neurons or changes in the structure of their correlations over time, we would expect to observe corresponding trends in their activation during the task or reactivation during sleep. We identified a subset of “drifting assemblies,” whose activation strength changed gradually across task and sleep sessions (Fig. 4a, Extended Data Fig. 11). Assembly drift was seen across assembly type and task/sleep epochs, but was greater during ripples (Extended Data Fig. 11, GLM β = 0.044, SE = 0.009, t(942) = 4.67, p < 0.001) and task (β = 0.024, SE = 0.009, t(942) = 2.65, p = 0.008) relative to the reference sleep-Nonripple epochs, and showed an interaction whereby Old assemblies exhibited less task drift (β = –0.023, SE = 0.011, p < 0.05, Extended Data Fig. 11).

**Fig 4.**
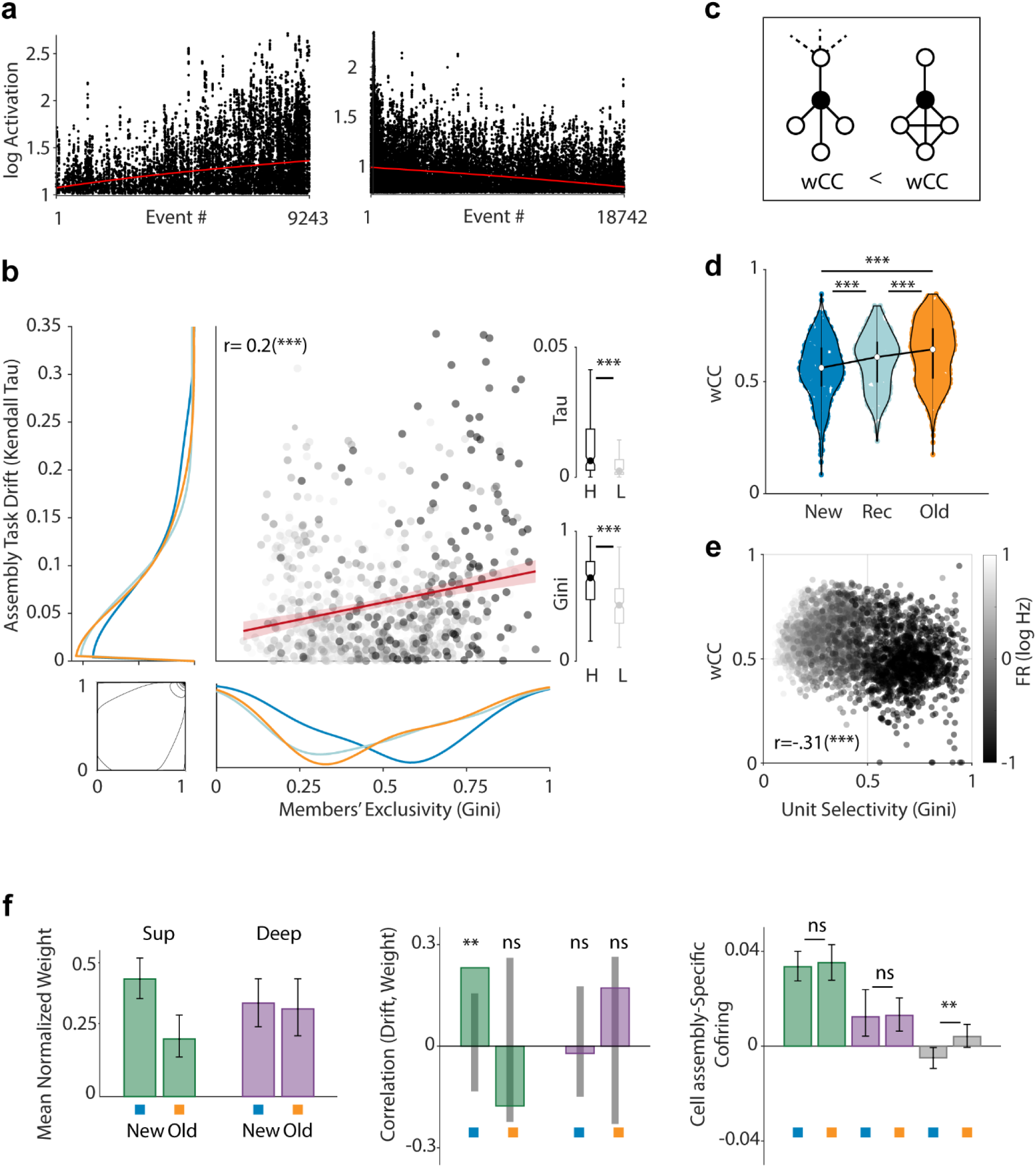
Network properties of assembly reorganization **(a)** Examples of drifting cell assemblies with positive (top) and negative (bottom) drift in activation strength across the task, indicated by red lines representing the Theil–Sen estimator. **(b)** Scatterplot showing the relationship between assembly drift during the task (Kendall tau) and the average exclusivity of an assembly’s members (Gini coefficient), shown with marginal distributions separated by memory category. Points are grayscale coded based on absolute changes in mean correlations from the first 25% to the last 25% of cell assembly activation events in the task (see methods for details). The red line represents the best-fit line with 95% confidence intervals, indicating a significant correlation from a permutation test (p < 0.001). The best-fitted copula model (Gumbel, θ = 1.20) is used to illustrate extreme joint positive dependencies, suggesting the relationship is pulled by the most extreme points (e.g. exclusive, drifting assemblies). (right) Distributions of Kendall Tau (top) and average member task participation Gini (bottom) for cell assemblies with the Highest 25% (black) and Lowest 25% (gray) largest changes in absolute correlation (*** p < 0.001, permutation test). **(c)** Schematic example illustrating two neurons with identical numbers of connections but different weighted-clustering coefficients (wCC), based on the connection strength among their neighboring neurons. The wCC incorporates edge weights to quantify the density and cohesiveness of local connections. **(d)** Violin plots depicting the distribution of wCC among assembly members across different categories (n = 568, 745, 1179 for new, recent, and old respectively). Pairwise differences from permutation tests with 10000 permutations and FDR correction. *** p < 0.001 **(e)** Scatterplot of wCC as a function of selectivity where each dot represents a neuron, grayscale-coded based on that neuron’s log10 firing rate (n = 3174). Spearman correlation (r); *** p < 0.001 permutation correlation test. **(f)** (left) Contribution of superficial (green) and deep (purple) pyramidal neurons to the formation of new (blue) and old (orange) cell assemblies, quantified as the mean normalized weight in PCA/ICA loadings (See Extended data fig. 12, GLM model: ‘Weight ∼ Age*Layer’ with significant main effect of the Age context (New/Old), and the Age:Layer interaction, p < 0.05). (middle) Correlation between mean normalized weight and Kendall tau, with 95% confidence intervals from a permutation test (**p < 0.01) (right) Cofiring between pairs of superficial (green), deep (purple), and superficial-deep (gray) member neurons within new (blue) and old (orange) cell assemblies (permutation test, ** p < 0.01).

Having observed assembly drift as a regular phenomenon, we evaluated whether drift might differ as a function of specific assembly characteristics. Assemblies’ constituent neurons can switch between assemblies and thus change their participation dynamics in a specific assembly’s activation^46^. A cell participating in many assemblies may be less able to maintain its strength among members of any one assembly, due to mechanisms such as heterosynaptic plasticity^47,48^. Accordingly, one might predict that assemblies whose members are more exclusive to that assembly could provide greater stability (less drift) than those with highly promiscuous members participating in competing activity patterns. To investigate if there is a relationship between the membership exclusivity of a cell assembly and its future drift, we computed average member participation Gini which is a measure of selectivity (see Extended Data Fig. 10 and methods for details) and correlated it with our measure of drift. New assemblies had higher Gini coefficients, suggesting increased selectivity in their members (Fig. 4b, pairwise permutation tests, 10000 permutations: New vs. Recent, *p* = 0.0026 and New vs. Remote, *p* = 6×10^-4^; Remote vs. Recent, *p* = 0.804). Furthermore, and in contrast to the predicted drift rates, assemblies with more exclusive members showed greater drift than those with more promiscuous members (Fig. 4b, r = 0.2, p < 0.001, Pearson’s correlation with permutation test).

These results align with our findings on cell assembly coupling, where both Old and Recent cell assemblies displayed greater inter-assembly correlation and inter-member cofiring. We also measured changes in pairwise cofiring across the initial and final 25% of activation events for each cell assembly and examined their relation to both membership exclusivity and drift. Cell assemblies exhibiting the largest drift and membership exclusivity also showed the greatest changes in cofiring (Fig. 4b). This suggests that assemblies initially isolated tend to drift toward greater integration within a larger, overlapping network. Observations in rodents indicate that drift results from alterations in higher-order network properties: richly connected neurons are less likely to be dropped from an assembly^42,49^. Here, the relation between experience (i.e. age of the assembly) and network connectedness was unclear, therefore we measured the clustering coefficient for members of cell assemblies to see if their constituents differed as a function of age category (Fig. 4c-d). The clustering coefficient reflects the extent to which a node’s neighbors form interconnected clusters, indicating local network density and structure (Fig. 4c). Assemblies showed larger weighted clustering coefficients (wCC) with increasing age (New < Recent < Old), suggesting increased interconnectedness and local cohesion, which may facilitate more stable and resilient network structures essential for long-term memory processing (Fig. 4d). Additionally, we observed a significant relationship between wCC and cell selectivity in specific assembly activations, further influenced by neuronal firing rates (Fig. 4e).

## Superficial and Deep CA1 cell participation across assembly age

Monkey CA1 pyramidal cells, like those in rat and mouse CA1, show a laminar segregation of function, including cell assembly membership^50^. In mice, the engagement and functional roles of superficial and deep pyramidal neurons depend on prior experience^51^ and the type of memory being processed^52^. To assess whether a substrata bias exists in monkey assembly participation as a function of experience, we calculated the mean normalized assembly weight of identified superficial and deep CA1 pyramidal neurons, indexing each cell’s contribution to cell assembly dynamics (Fig. 4f and Extended Data Fig. 12). Superficial pyramidal cells preferentially participated in New assemblies, whereas deep pyramidal cells did not (Fig. 4f), similar to observations in rodent CA1^51^. In addition, we found a significant correlation between the degree of superficial cell participation in an assembly and the assembly’s future drift rate (Fig. 4f), potentially driving the faster drift rates seen in New assemblies. Furthermore, the layer segregation of assemblies was not constant across experience. With assembly age, for those superficial cells remaining, there was greater cofiring with deep cells than that seen for the New assemblies (Fig. 4f). Intra-sublayer cofiring, in contrast, remained unchanged as a function of age. Together, these findings suggest that superficial pyramidal cells initially contribute more to New assemblies, signaling future drift, and that, over the course of experience, more cross-strata interactions emerge.

## Discussion

This study provides insights into how the primate brain flexibly and dynamically reorganizes as a function of the individual’s history of learned experiences. We recorded from ensembles of single units in freely moving macaques during a memory-guided item-sequence task and in subsequent sleep, revealing the activation and reactivation of hippocampal and extrahippocampal cell assemblies associated with newly, recently, and remotely acquired experiences. We found that sleep reactivation is shaped by prior experience, with older and recently formed memories evoking stronger and more stable assembly reactivation compared to newly learned sequences. In contrast, cell assemblies associated with novel experiences underwent flexible reorganization and gradually formed higher-order links with assemblies representing older memories, giving rise to ‘metassemblies’—hierarchically coordinated networks that may support the integration of new and old experiences. We found that assemblies composed of more exclusive members were less stable over time (i.e. showed greater drift), suggesting that memory systems favor integration into a broader network rather than maintaining separability through sparse representations. To further understand the cellular basis of these dynamics, we examined the laminar organization of CA1 and found that superficial pyramidal neurons disproportionately contributed to new-biased assemblies and were associated with greater drift, whereas deep pyramidal neurons were more consistent across time. As memories aged, coactivity between superficial and deep pyramidal cells increased, suggesting that, like the metassembly results, subpopulations become progressively intertwined to support established memories. Collectively, these findings highlight flexible and experience-dependent organization of memory-related neural ensembles and identify network-level features that may underlie long-term memory formation and integration in primates.

Replay—the reinstatement of coordinated cell assembly patterns that were originally expressed during waking experience—is widely regarded as a key systems-level mechanism for memory consolidation, particularly during offline states such as sleep^53–55^. Much of this understanding is based on rodent studies, and evidence for analogous cell assembly dynamics and reactivation processes in primates remains comparatively limited. In monkeys, experience-related reactivation in neural ensembles was first observed during daytime rest and pauses in exploration or task performance^9^. In humans, similar reactivation patterns in lateral and medial temporal lobe ensembles have been linked to explicit memory retrieval^11,12,56,57^ and in the human and non-human primate motor cortex during post-learning naps^13,58,59^. Our present results extend these findings by demonstrating that, in monkeys, discriminable assemblies observed during extended episodic experiences reactivate during overnight sleep. Old– and recent-biased assemblies reactivated with greater strength during task and sleep than New-biased assemblies. Previous studies of reactivation or replay have not compared memoranda that differ in initial experiences with lapses extending over weeks; however, when mazes differ in familiarity, reactivation and replay magnitudes have been reported to favor novelty^60^, familiarity^22,36^, or other factors such as recent repetition and reward^61 37^.

We found in our superficial CA1 pyramidal cells a greater tendency to participate in new-biased cell assemblies, and their contribution to assembly dynamics predicted the assembly’s future drift rate. This finding echoes mouse studies showing greater long-term stability among deep-layer pyramidal neurons, and that superficial neurons exhibit activity changes between novel and familiar environments, whereas deep neurons show no such preference^51^. This may reflect differential synaptic plasticity: in mice, superficial neurons become more potentiated at CA3 synapses following experience, resulting in increased participation and elevated firing rates during post-experience sharp-wave ripples^62^. In contrast, whereas in vivo activity depresses CA3 input to deep pyramidal neurons, it does not significantly alter their firing rates across all SWRs. Furthermore, pyramidal cell–interneuron interactions also vary as a function of strata. Interneurons can actively participate in the formation of memory assemblies by modulating their coupling with pyramidal cells during learning, and such interactions persist into sleep^63^, thus the strata effects may be mediated by their interactions with inhibitory cell populations^50^. Previously, we showed that superficial and deep pyramidal cells exhibit strata-specific spike-timing interactions with inhibitory interneuron groups and tend to form non-overlapping cell assemblies, suggesting parallel information processing channels in CA1^50^. A more detailed analysis in the current study revealed that superficial and deep pyramidal cells that were initially negatively correlated became increasingly positively correlated during the processing of established old memories. In mice, distinct pyramidal subpopulations have been shown to encode different aspects of task structure—such as context, choice, and reward configuration^64–66^. Thus, the increased correlation between superficial and deep cells in our study may reflect a strengthening association between contextual cues, correct choices, reward outcomes, and affordances embedded in the task environment. A nonmutually exclusive possibility is a progressive recruitment of superficial cells into broader hippocampal cell assemblies that may serve to stabilize memory representations and enhance robustness—though potentially at the cost of increased mnemonic rigidity^52^. These findings indicate that the rates and structure of cell assembly reorganization may be constrained by CA1 microstructure^67^, irrespective of species.

Assemblies differed not only in their reactivation according to their age categories, but also in their interactions with other assemblies. During the task, a given assembly showed a wide range of preferences to co-activate with (or avoid activating with) other assemblies (i.e. ‘metassemblies’), regardless of whether the assembly pairs were new, old, or mixed. In sleep, however, those pairwise preferences from the task were preserved only when the metassemblies included older (recent or old) assemblies. One possible advantage to this higher-order structure is that these interactions support the representation of extended experiences^38,68,69^ and enable the integration of distinct memories separated by time^17,70^. Additionally, according to the “information overlap to abstract” theory, higher-order recombination of memory elements facilitates schema formation by selectively reinforcing shared features across related experiences^71–73^. This mechanism is critical for generating abstract, generalized knowledge^74,75^. Supporting this, empirical findings show that by flexibly linking smaller neuronal subpopulations into longer sequences, the hippocampus can spontaneously generate internal predictive models used in future experiences^76^. Once such a model is established, the memory system can bias replay content toward elements consistent across experiences—effectively sampling from an internalized model of the environment^77^. In this view, existing schemas not only shape how new memories are integrated but also determine which are prioritized for consolidation. Consistent with this, the content of awake sharp-wave ripples (SWRs) has been proposed to serve as a neurophysiological tagging mechanism, to selectively preserve and consolidate aspects of experience for future use^40^. In light of the positioning of SWRs during waking recall of multiple experiences in primates^20,21,78,79^, such a mechanism could extend the role of replay beyond consolidating specific episodes, to include a role in extracting and reinforcing generalizable structure.

## Acknowledgments

**Funding:** The funding for this study was provided by National Institute of Neurological Disorders and Stroke grant (R01) 1RF1NS127128-01, NEI (P30 EY008126), NIH OD (1S10D021771) to the VUIIS Center for Human Imaging, and the Whitehall Foundation grant 100001391. **Author contributions:** <u>Conceptualization</u>: S.A, and K.L.H; <u>Data curation</u>: S.A, and A.A.; <u>Formal analysis</u>: S.A, and K.L.H; <u>Funding acquisition</u>: K.L.H; <u>Investigation</u>: S.A, and K.L.H; <u>Methodology</u>: S.A, and K.L.H; <u>Software</u>: S.A, A.A.; <u>Supervision</u>: K.L.H; <u>Visualization</u>: S.A, A.A., K.L.H; <u>Writing – original</u> <u>draft</u>: S.A, and K.L.H; <u>Writing – review & editing</u>: S.A, and K.L.H; **Data and materials availability**: All data are available in the manuscript or the supplementary materials.

## Extended figures

**Extended Data Fig. 1:**
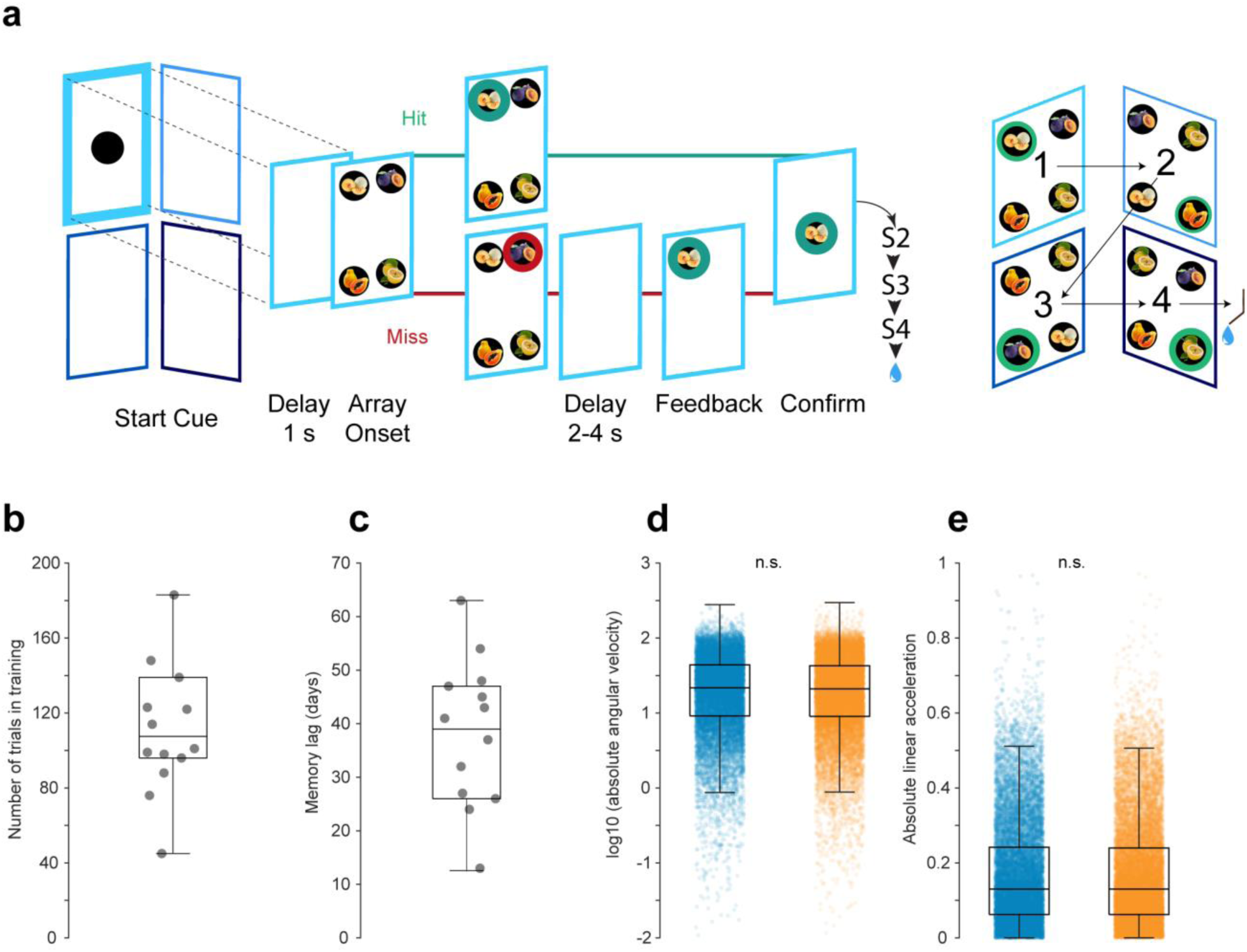
Sequential item-in-context association task **(a)** Left: Trial sequence for the first screen in the task sequence. The bold outline represents the active screen; the green ring indicates the correct target, and the red ring indicates an incorrect distractor. After completing this procedure on one screen, the trial proceeds to the next screen in the sequence finishing with reward, which followed every 4^th^ screen completion. Right: Four objects and example positions on the screen for one trial. Green rings indicate the associated target objects assigned to each screen. The assignment of objects to the four screen positions in the 2 × 2 array is randomized across screens and trials, preventing prediction of object location, and the background image is simplified for illustration purposes. **(b)** Distribution of the number of training trials prior to the first day of recall for the old set. **(c)** Distribution of memory lag. Number of intervening days between the end of initial exposure to a stimulus set during learning and its return as the ‘Old’ category. **(d)** Distribution of the angular velocity values (degrees per second) during trial epochs (n.s., permutation test). **(e)** Distribution of the linear acceleration values (m^2^/s) during trial epochs (n.s., permutation test). New set: Blue, Old set: Orange.

**Extended Data Fig. 2:**
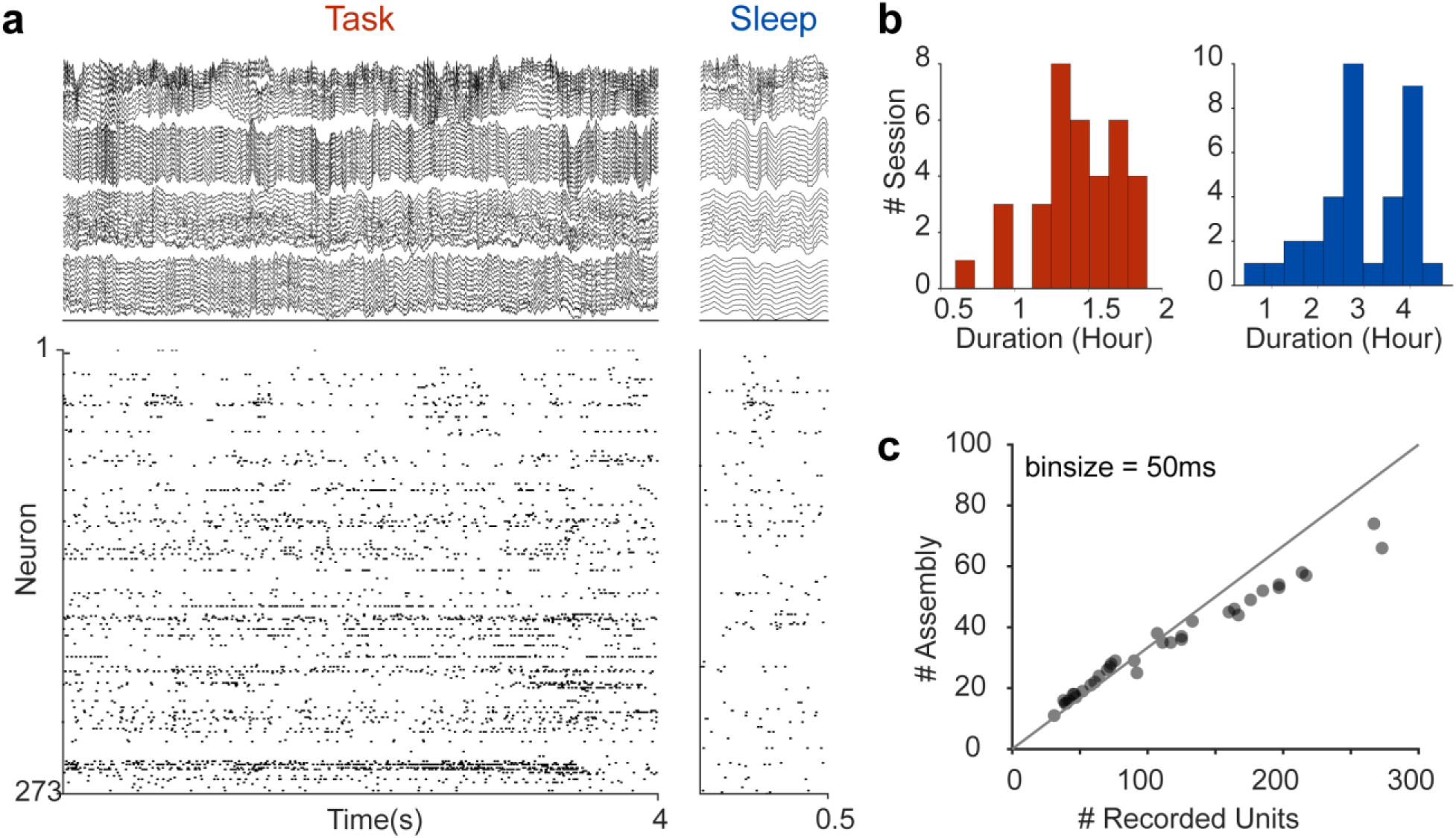
High-density ensemble recording in freely moving macaques during task and sleep **(a)** High-density wireless recordings showing a subset of local field potentials (top) and ensemble spiking activity (bottom) during task performance (left) and post-task sleep (right). The example sleep epoch is aligned to a detected sharp-wave ripple event. **(b)** Distribution of wireless recording durations during task performance (red, left) and sleep (blue, right). **(c)** Relationship between the number of recorded units in a session and the number of detected assemblies using the PCA/ICA method.

**Extended Data Fig. 3:**
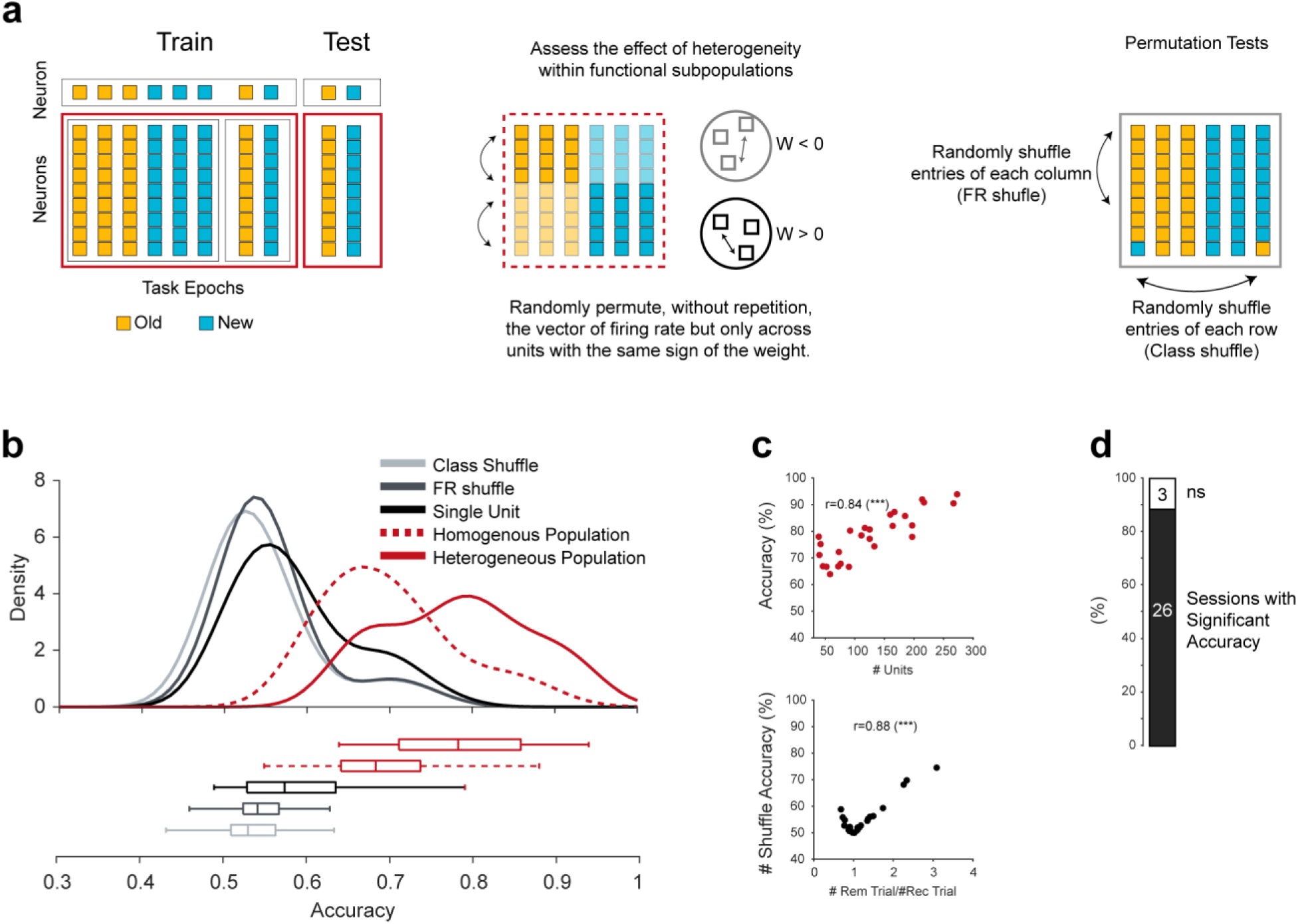
Decoding behavioral context from population spiking activity **(a)** Schematic representation of the linear support vector machine (SVM) training and testing approach. Left: Main (heterogeneous) population decoding and single-unit decoding. Middle: Homogeneous population decoding. Right: Class and firing rate shuffling tests for significance (see Methods and Ref ^80^ for details). **(b)** Kernel density distributions of SVM cross-validation accuracies for different design matrices. **(c)** Top: Correlation between the number of units included in population decoding and SVM performance accuracy. Bottom: Correlation between trial number balance across conditions and the accuracy of the shuffled design matrix. **(d)** Proportion of sessions with significant decoder accuracy.

**Extended Data Fig. 4:**
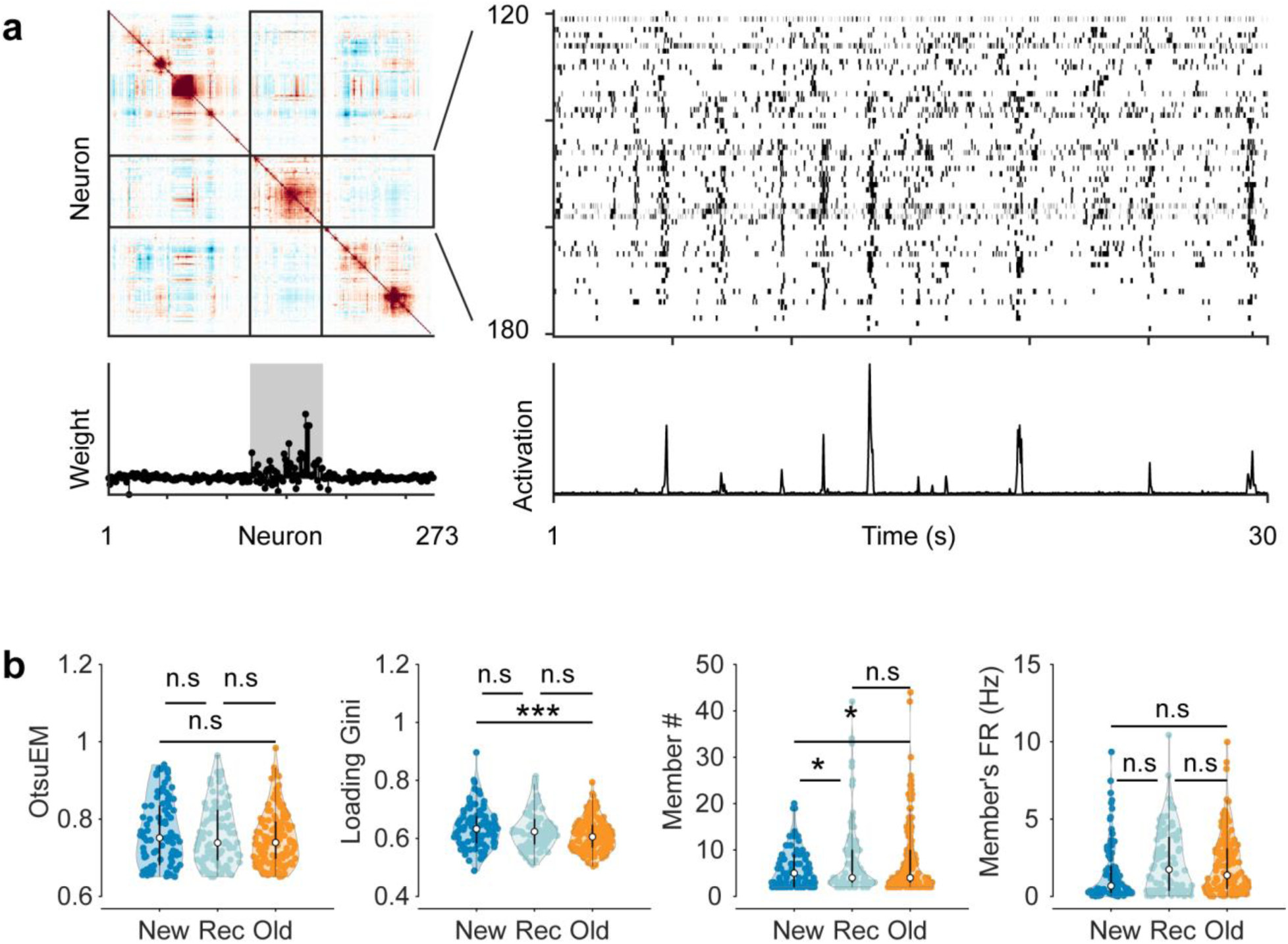
Cell assembly detection and quality control **(a)** Top-left: Correlation matrix (r) showing pairwise correlations between units in an example session. Top-right: Firing rate traces of a subset of neurons exhibiting highly correlated activity. Bottom-left: Example cell assembly pattern (1 of 66 detected during the session), with significant cell assembly members highlighted in gray. Bottom-right: Activation strength trace of the highlighted cell assembly and the corresponding firing rate raster. **(b)** Left to right: Distributions of OtsuEM values, loading Gini coefficients, number of members, and members’ firing rates for new-, recent-, and old-biased cell assemblies.

**Extended Data Fig. 5:**
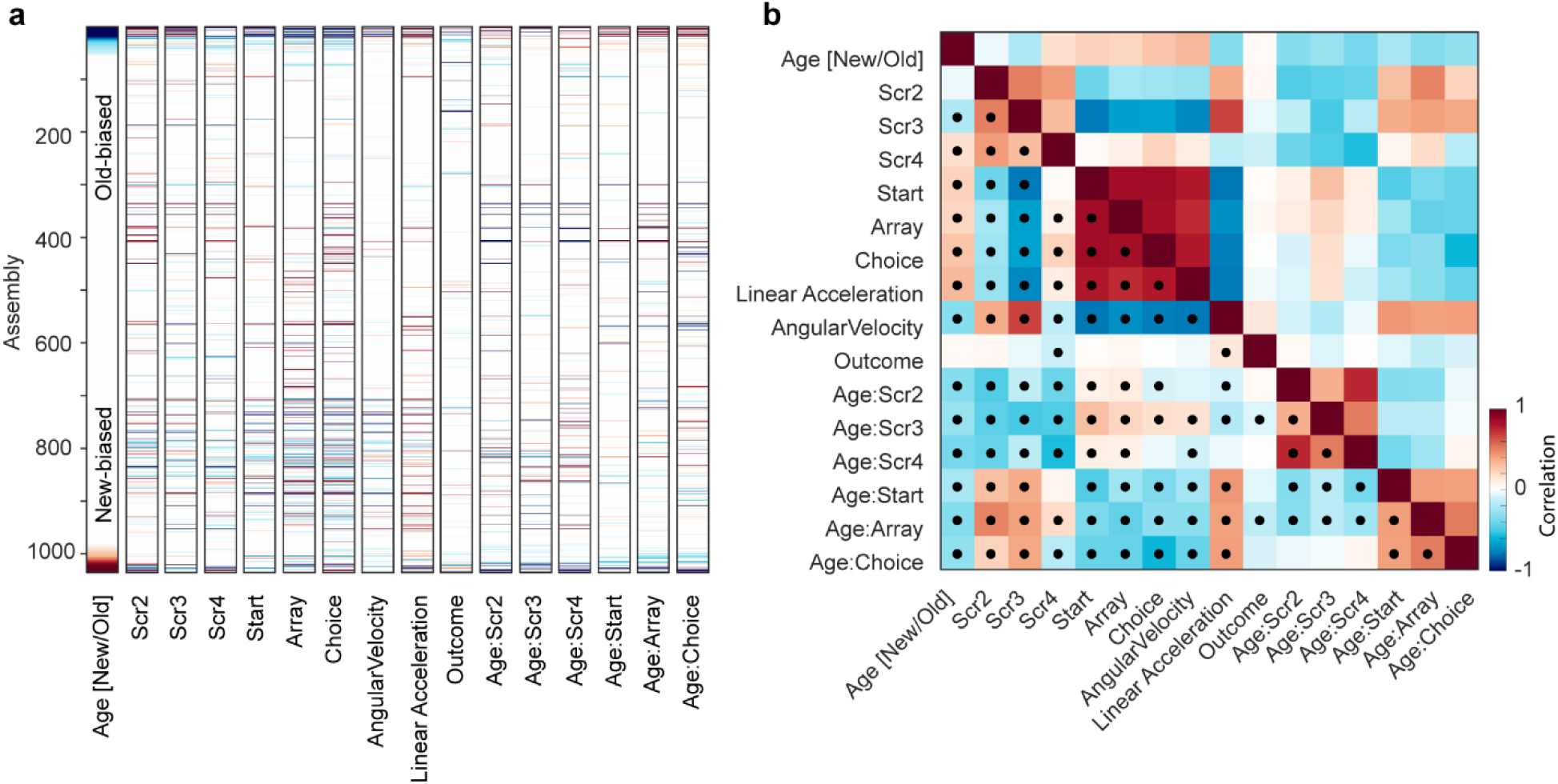
Cell assembly activations are selectively modulated by diverse task variables. **(a)** Distribution of the beta coefficient weights for the generalized linear model. Nonsignificant beta coefficients (p > 0.05) were set to 0 for the visualization. Beta coefficients are sorted based on the age column. **(b)** Pairwise correlations of the beta coefficients vectors (in a.) showing the task-related covariates across the population of cell assembly activations. Dots indicate significant correlation values (p < 0.05, permutation test with FDR correction).

**Extended Data Fig. 6:**
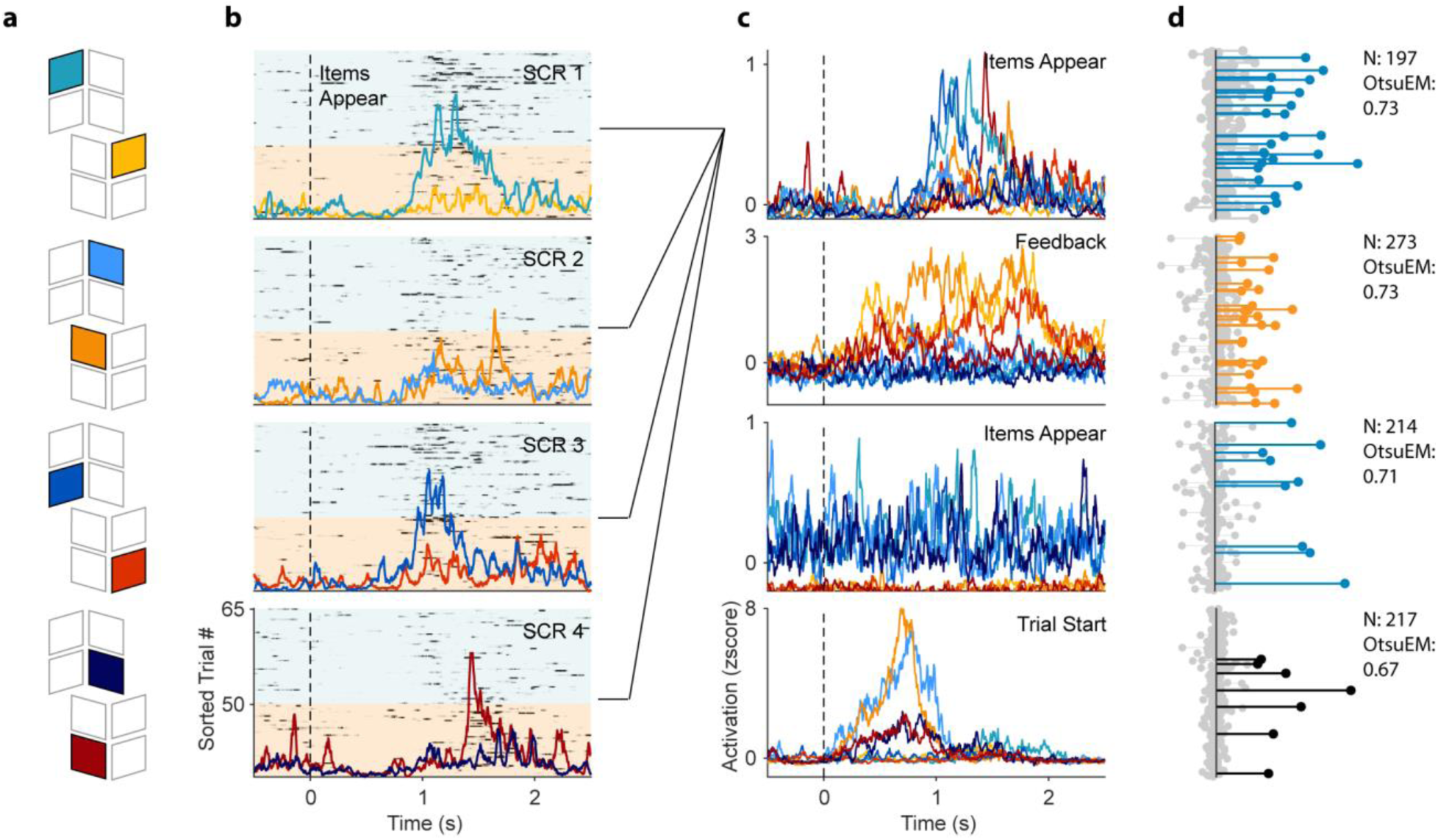
Context-biased cell assembly activation during task. **(a)** Schematic of screen locations in the 3D task environment “Treehouse.” **(b)** Activation strength across trials and screens for an example assembly, along with its average activation per screen. Trials are sorted by context for clarity (New: blue; Old: orange). Average normalized activation strengths are color-coded by the screen location during each trial. **(c)** Examples of context-biased and/or task-selective activations during different task epochs. Each plot shows the average activation trace of a different cell assembly across screen locations within a given context. Traces are color-coded by screen location. The top plot is the assembly example shown in (a). **(d)** Assembly templates (loadings) corresponding to the assemblies shown in (c). Significant weights are color-coded by bias (New: blue; Old: orange; Nonbiased: black). *N* indicates the number of neurons included in the PCA/ICA analysis. *OtsuEM* represents the effectiveness measure of Otsu thresholding. Only high-quality assemblies with an OtsuEM > 0.65 were included in the biased cell assembly analysis.

**Extended Data Fig. 7:**
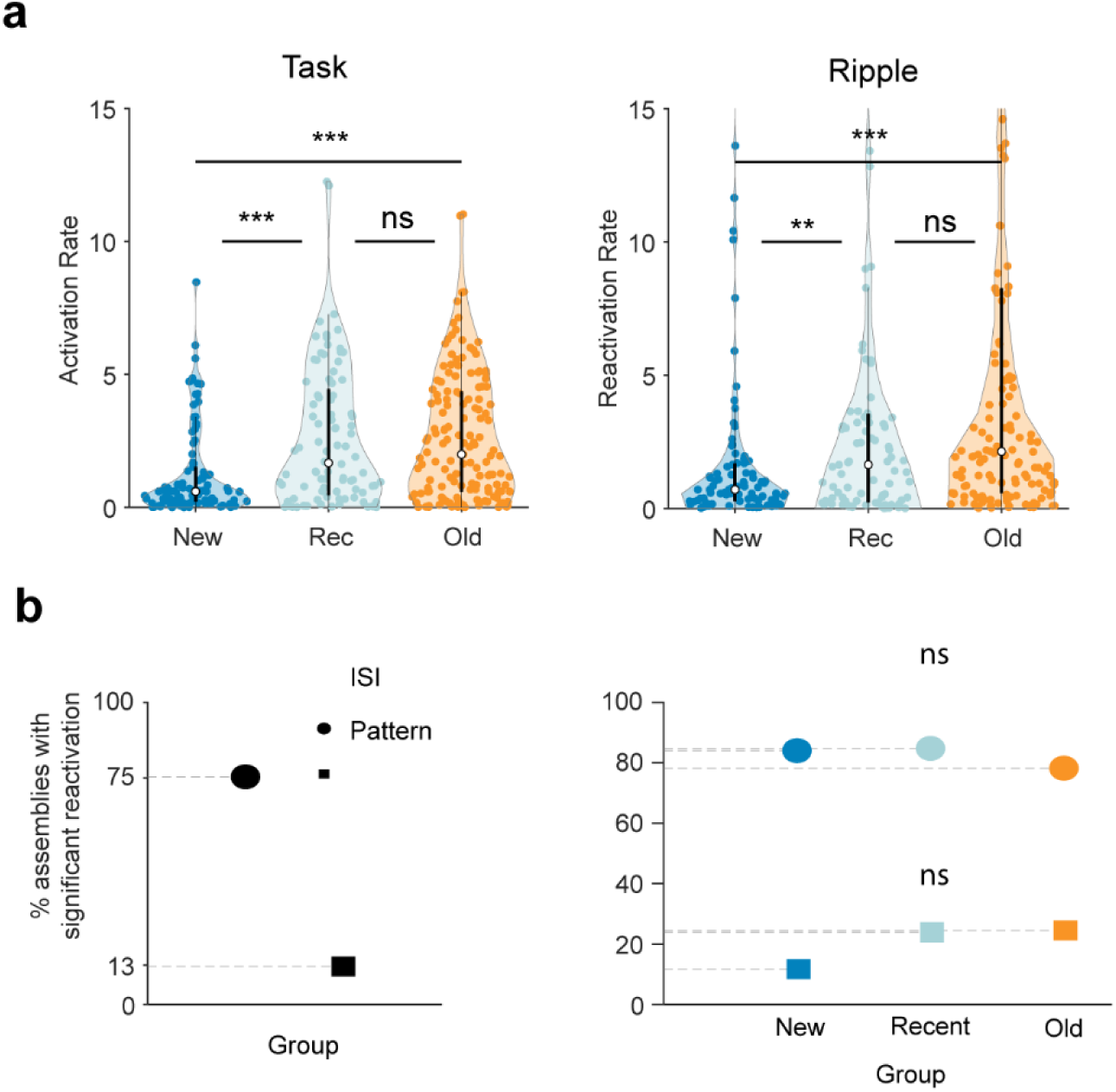
Cell assembly dynamics across different behavioral states **(a)** Activation and reactivation rate during task performance and sleep ripple for different categories of cell assemblies (*** p < 0.001, pairwise permutation test with FDR correction). **(b)** Left: Proportion of cell assemblies with significant reactivation during sleep, based on two different significance measures (see Methods). Right: Proportion of reactivating cell assemblies separated by new-, recent-, and old-biased categories.

**Extended Data Fig. 8:**
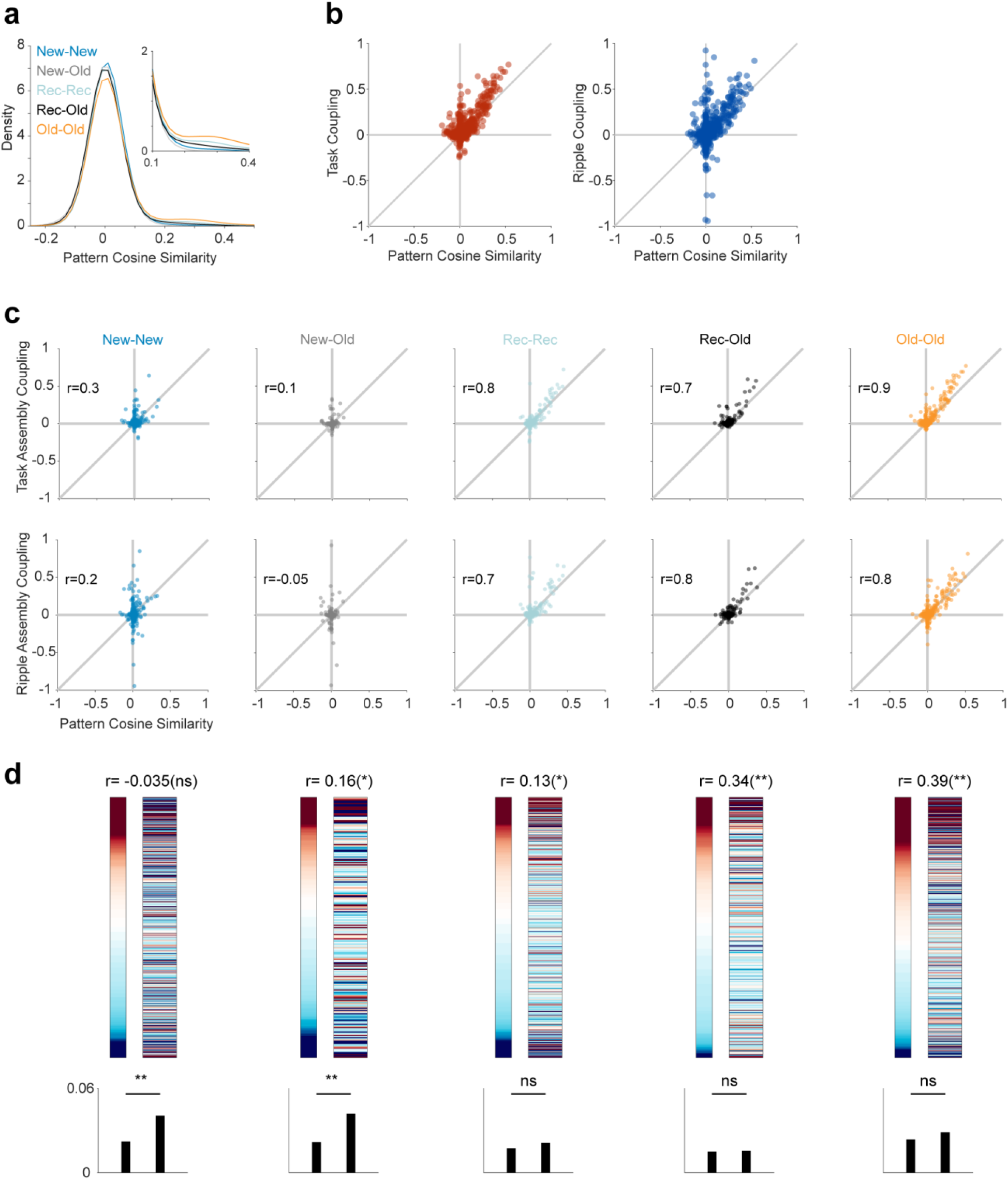
Pattern similarity and cell assembly coupling **(a)** Distribution of pairwise pattern cosine similarity between different categories of cell assemblies. **(b)** Relationship between pattern cosine similarity and cell assembly coupling across various brain-behavioral states. **(c)** Assembly coupling during task (top) and sleep ripple (bottom) as a function of pattern cosine similarity, separated by cell assembly category. **(d)** Same as Figure 3, but calculated exclusively for pairs of cell assemblies with cosine similarity < 0.15.

**Extended Data Fig. 9:**
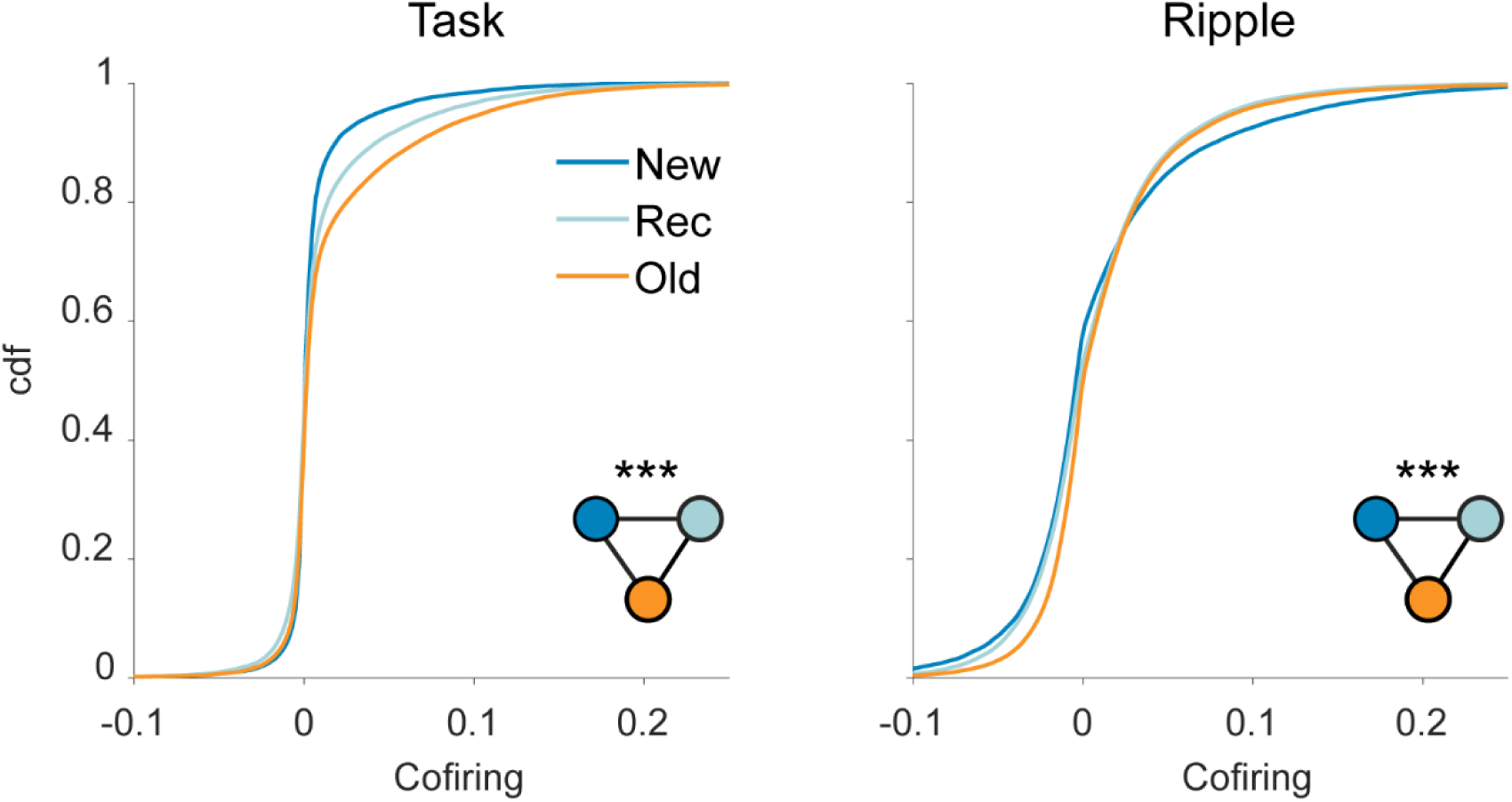
Cofiring dynamics among cell assembly members across brain-behavioral states Cumulative density function of global cofiring among members of cell assemblies with different categorical biases across various brain-behavioral states. Inset: *** p < 0.001 pairwise distribution differences indicated by black lines connecting color-coded circles, permutation test)

**Extended Data Fig. 10:**
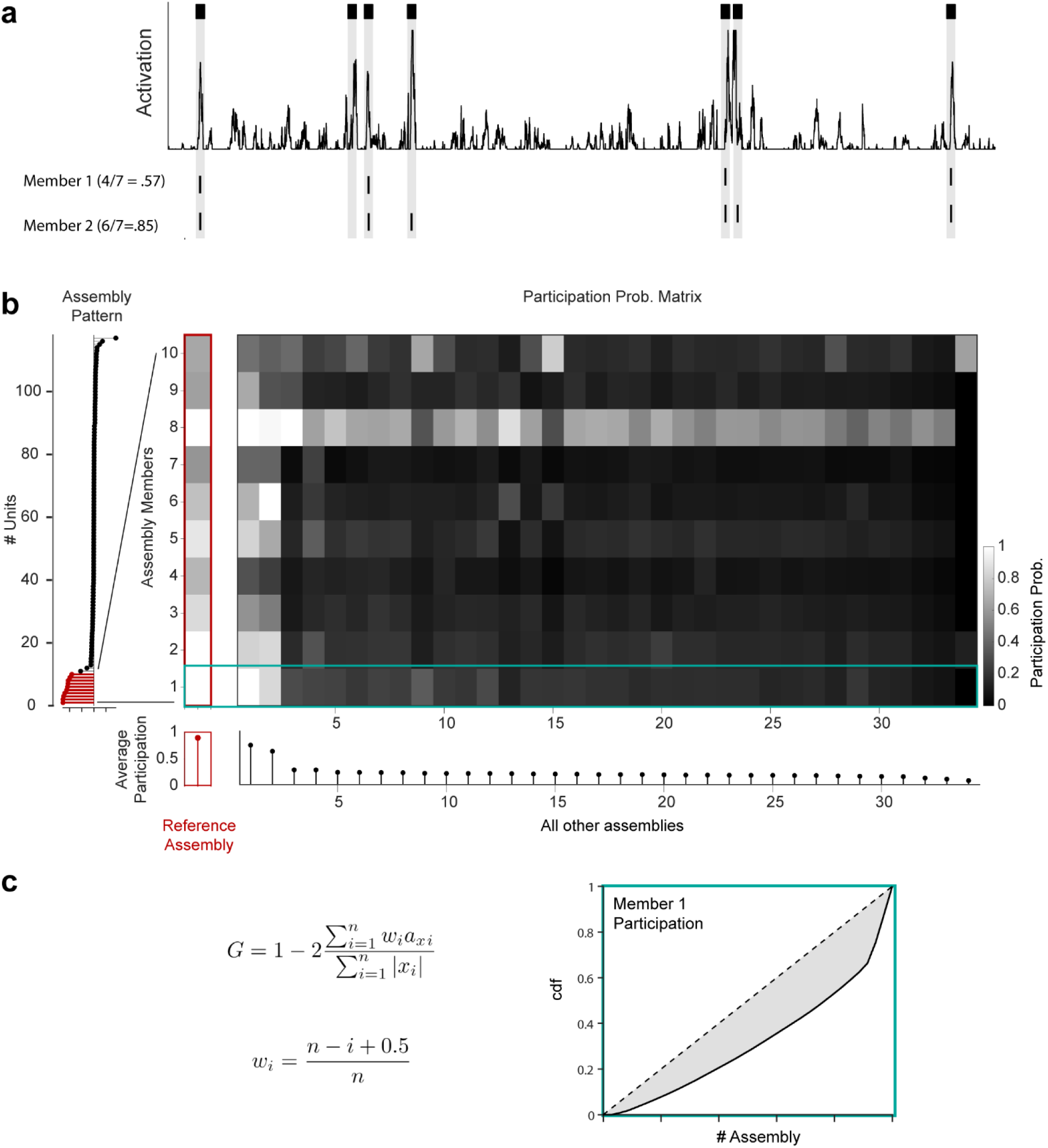
Schematic representation of assembly exclusivity and member selectivity quantification **(a)** Activation trace plot for a single cell assembly alongside the spiking activity of various member neurons. The participation probability of a neuron in a specific cell assembly was calculated as the number of activation events where the neuron fired at least one spike, divided by the total number of detected activation events. **(b)** Color plot depicting the participation probability matrix for members of a reference cell assembly (left) during activation events of all detected cell assemblies within a session. The bottom plot shows the average participation probability of the target members during each detected cell assembly in the session. Higher exclusivity of a reference cell assembly is indicated by lower average member participation probabilities across all but the reference cell assembly. **(c)** A selectivity index was calculated for each member based on the distribution of participation probabilities across all detected cell assemblies. The selectivity index was determined as the Gini coefficient of the participation probabilities (see Methods section for details).

**Extended Data Fig. 11:**
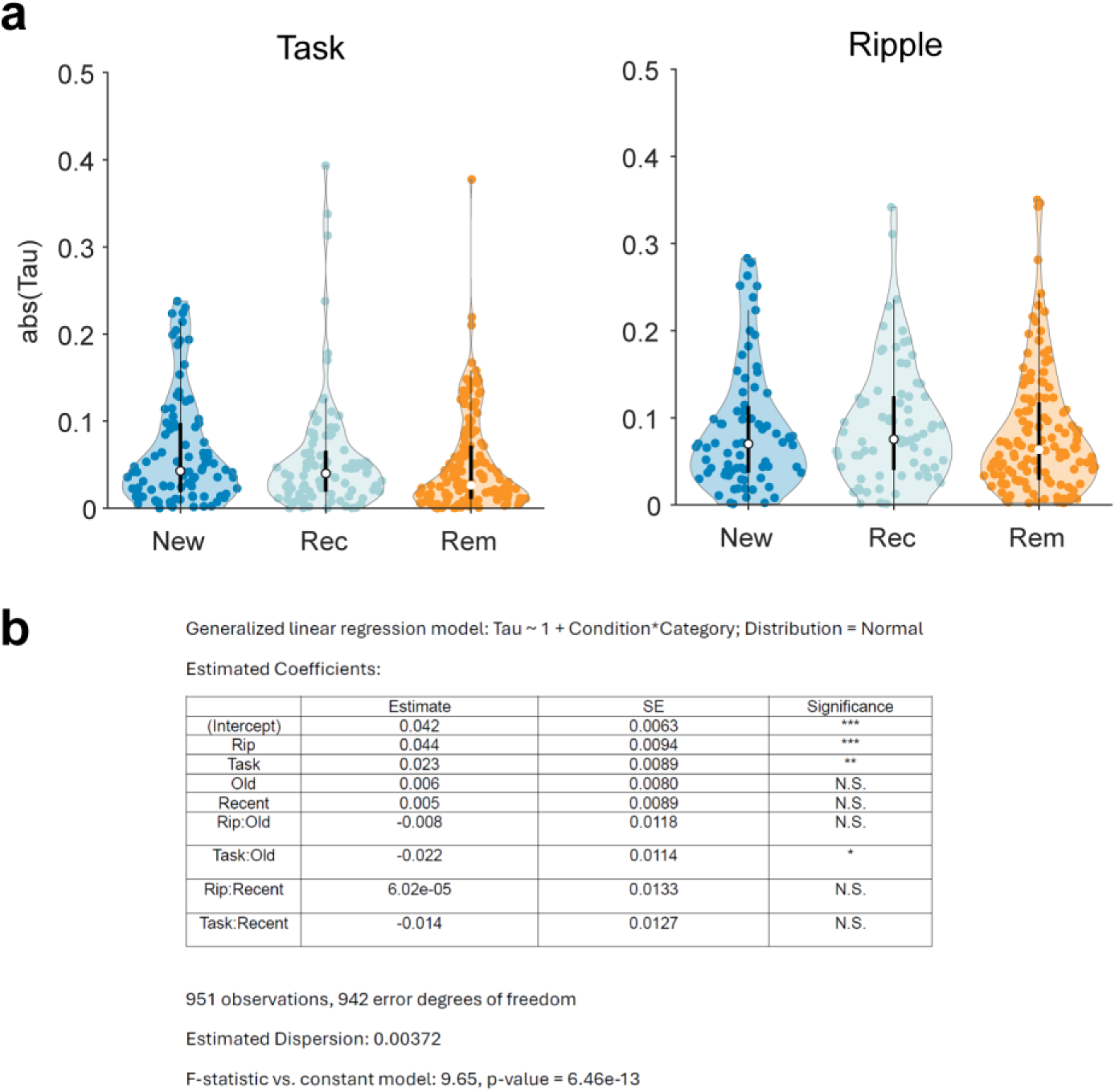
Drift dynamics across behavioral states **(a)** Distribution of absolute Kendall’s tau values for different categories of cell assemblies across brain-behavioral states. **(b)** Generalized linear model (GLM) results predicting drift (Kendall’s tau) using the cell assembly features: state (Ripple, Task) and sequence age type (Recent, Old).

**Extended Data Fig. 12:**
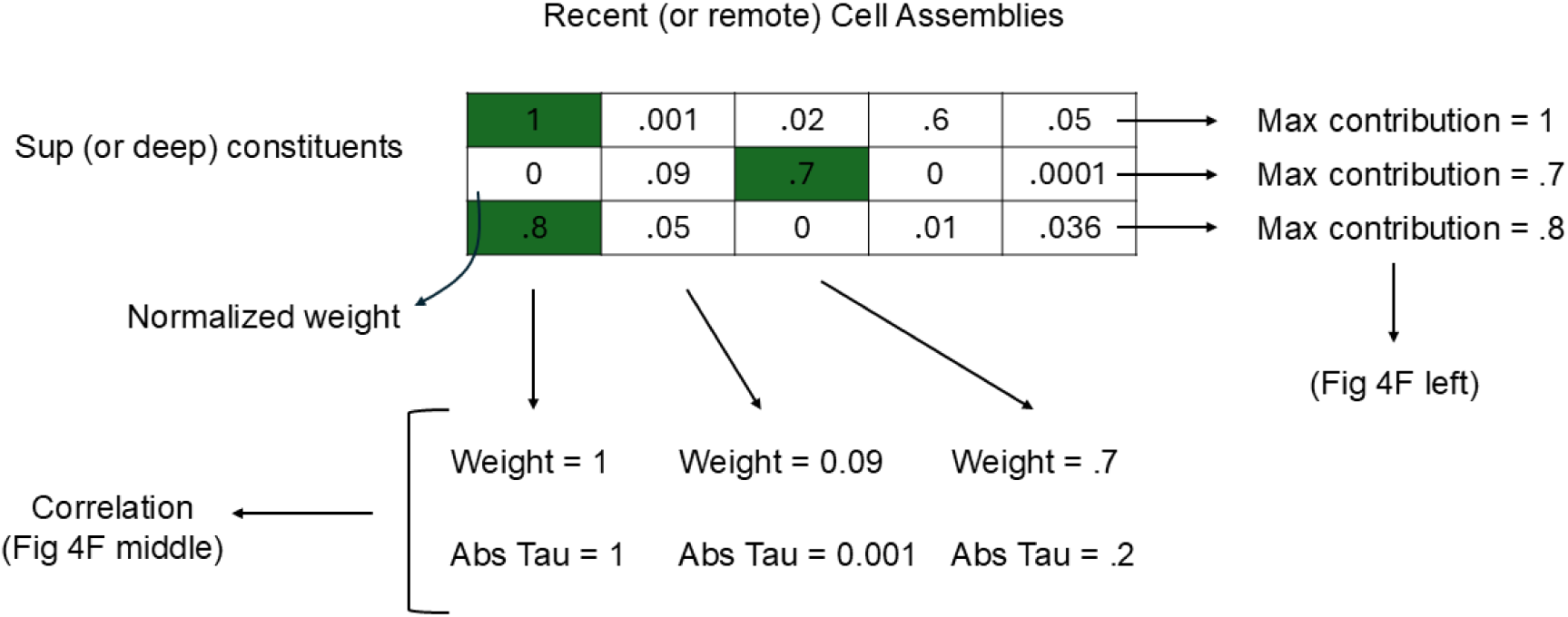
Superficial-Deep Contribution to New and Old Cell Assemblies (see Materials and Methods for full description).

## Materials and Methods

### Subjects and Behavioral Design

Two adult female macaques (*Macaca mulatta*, referred to as ‘M1’ and ‘M2’) were subjects in this study. All animal procedures were approved by the Vanderbilt Institutional Animal Care and Use Committee, in compliance with the policies of the United States Department of Agriculture and Public Health Service on the humane care and use of laboratory animals. Details of the testing enclosure, task, and training were previously described in Ref ^28^.

#### Trial Design

Both monkeys underwent training in a 3D testing enclosure, which allowed them to move freely. This enclosure was equipped with eight touchscreens distributed around its periphery (Fig. 1 and Extended data fig. 1). The task started with the presentation of a start cue (black circle) that designated the active screen for the animal. Upon touching the cue, and after a 1 s delay, the array onset consisted of the appearance of 4 objects presented in a 2 by 2 grid on a background scene providing context that was specific for that set of objects. Selection: If monkeys touched the correct item (Target’) – the item associated with that screen – a ‘correct’ tone was played as the four objects disappeared. If, instead, they selected one of the 3 non-target objects (distractors’), an ‘error’ tone was played along with the disappearance of the stimuli for 2-4 seconds, and then, the correct target appeared in place. Confirmation: in both cases, before proceeding to the next screen in the sequence (or, after screen 4, reward delivery), the target reappeared at the center of the screen and monkeys had to touch it. Subjects proceeded through the same trial sequence across all 4 screens, in order, before receiving fluid reward in the opposite corner of the treehouse. For M1, the total reward was calculated based on the performance of the animal within a trial (e.g., 2 drops of juice per correct selection and 0 drops for incorrect touch) and delivered at the reward receptacle. For M2, a fixed reward amount was delivered, but the reward type (flavor) was switched in the middle of the session to maintain motivation. For the presented data, both subjects always traversed an identical sequence starting with screen 1 (upper left) to screen 4 (lower right) on one corner of the environment.

#### Session design

During each session, monkeys completed various trial blocks distributed between the new and old conditions. These conditions shared the same trial structure but were situated in opposite corners of the testing apparatus. New trials (day 1) were composed of item-context stimuli that monkeys had never seen previously and had to learn the associations. Monkeys followed up their training and further learning of the new associations on day 2 (recent trials). Old trials were composed of item-context stimuli that monkeys learned (i.e. they had been New sets) 2-9-weeks prior to testing with no intervening exposure to them until recall. For Old trials, we concatenated Day 1 and Day 2. Old/New set assignments are balanced across corners, because each New set that is assigned to the opposite corner from the paired Old set, eventually becomes and Old set itself.

#### Sleep

Following their training session, monkeys were returned to their housing. For monkey M2, overnight recordings started within approximately 20 minutes of the training session. For monkey M1, they occasionally included 1–2-hour gaps. Sleep epochs occurred with the monkeys in their normal housing area, with their usual social housing accommodations, in complete darkness, following the automated lighting system’s 12/12 light/dark cycle. Throughout the manuscript, we used the terms “task” and “sleep” to broadly refer to recordings in the 3D testing enclosure and overnight occupancy in the housing, respectively.

### Electrophysiological recordings

Active multichannel probes were mounted on microdrives and inserted into a chronically implanted base ^81^. These included a 128-channel probe (DA128-1, linear configuration with 40μm contact spacing) for Monkey M1 and four 64-channel probes (custom design, organized into 4 parallel shanks with 40 channels at 90 μm spacing and 3 shanks with 8 channels at 60 μm spacing) for Monkey M2 (’Deep Array Probes’, by Diagnostic Biochips, Inc., currently distributed through Cambridge Neurotech, Inc.). The microdrives (M2: NanoDrives, Cambridge Neurotech, Inc.; M1: custom, Rogue Research, Inc.) facilitate precise depth positioning adjustments post-implantation, and allowing the raising and relowering (max. 7 mm or 5 mm, respectively) into target areas while remaining implanted. Post-operatively, the probes were typically advanced in 125 μm steps until the target positions were achieved. For M1 the probe sampled from anterior hippocampal CA1. For M2, two of the probes sampled from anterior CA1, and one sampled the posterior hippocampus. In both monkeys the remaining probe targeted areas in and around the retrosplenial and posteriomedial cortex. As previously described ^50^, The localization of recording sites was verified through postoperative CT scans, coregistered with pre-operative MRI data, and also by referencing functional landmarks that changed with increasing depth (for hippocampal probes). Notably, the emergence of depth-specific sharp-wave ripples (SWRs) within unit-dense layers served as a key reference point for CA1. Local field potentials (LFPs) were digitally sampled at a rate of 30 kHz using the FreeLynx Wireless Acquisition system (Neuralynx, Inc.) and were subsequently bandpass filtered within the 0.1 Hz to 7500 Hz range. During task performance and sleep recordings for Monkey M2, data were wirelessly transmitted to the Freelynx acquisition system (Neuralynx, Inc.). To optimize battery life, sleep recordings for Monkey M1 were stored on an SD card within the acquisition system. A high-frequency noise signal at approximately 6 kHz appeared at low (∼10%) battery capacity. The onset of this noise was detected by calculating the root mean square (RMS) envelope of the band-passed filtered signal between 5800-6200 Hz. The noise initiation point was defined as the timestamp at which the RMS envelope exceeded 110% of the median value, and sleep data following this point was excluded. For the purpose of isolating stable single units across task and sleep epochs, data were initially converted to microvolt units, bitVolts were standardized to 0.195, and the data were subsequently transformed into binary files encoded as 16-bit integers. Only sessions that exhibited no signal loss during recording were included in the analyses. In LFP-related analyses, the raw signal was subjected to third-order Butterworth filtering with a low-pass cutoff frequency set at either 350 Hz or 450 Hz, and the data was downsampled to 1 kHz. Portions of the data from the CA1-localized probes have been published previously ^50^.

### CT-MRI image processing and coregistration

The General Registration tool, Elastix, in the Slicer (version 4.11), was used to perform the registration of post-operative CT scans with pre-operative MRI images. The registration process maintained default parameters. Preceding registration, the CT images were cropped near to and including the cranium.

### Inertial Measurement Unit (IMU) Data Collection and Processing

Inertial data were acquired using a triaxial accelerometer embedded in the Freelynx system, sampled at 3 kHz along the X, Y, and Z axes. Linear acceleration (BASE_LAX, BASE_LAY, BASE_LAZ) and angular velocity (BASE_AVX, BASE_AVY, BASE_AVZ) signals were collected and z-scored. Each signal type was concatenated into a 3-dimensional matrix and subjected to Principal Component Analysis (PCA) to reduce dimensionality and identify the dominant axis of variance. The first principal component (PC1) from each PCA was extracted and used in subsequent analyses.

### Detecting hippocampal sharp-wave ripples and estimating ripple slope

A single hippocampal LFP channel with largest ripple amplitude was selected for ripple detection. The wide-band signal was band-passed filtered between 100-180 Hz using a 3^rd^ order Butterworth filter, and squared signal was calculated. The squared signal was further band-passed filtered in 1-20 Hz range and z-normalized. SWR peaks were detected by thresholding the normalized squared signal at 3 SDs above the mean, and the surrounding SWR start and stop times were identified as crossings of 1 SDs around this peak. SWR duration limits were set to be between 40 and 400 ms. Adjacent events with an inter-ripple interval of < 20 ms were merged.

#### Exclusion Criteria

In addition to assessing amplitude and duration, we used several criteria to identify and exclude potential false-positive ripple events.

#### Channel Noise Exclusion

A ‘noise’ channel, defined as one devoid of detectable sharp-wave ripples (SWRs) in the local field potential (LFP), was designated. Any events simultaneously detected on this channel were considered as potential false-positives, likely originating from artifacts such as electromyography^82^.

#### Spectral Analysis

We conducted spectral analysis of each detected ripple event to further scrutinize its characteristics. The data was spectrally decomposed using Morlet wavelets, allowing us to compute the frequency spectrum for each event. This was achieved by averaging the normalized instantaneous amplitude within ±50 ms of the ripple peak over the frequency range of 50-200 Hz, normalized by multiplying the amplitude by the frequency. We then analyzed the number and properties of spectral peaks in each detected ripple frequency spectrum. These peaks were identified using the findpeaks function in MATLAB, considering parameters such as peak height, prominence, peak frequency, and peak width. This analysis aimed to ensure that the detected ripple events genuinely reflected high-frequency, narrowband bursts within the ripple band range. We applied multiple criteria to achieve this: first, authentic ripple events were expected to exhibit a predominant spectral peak within the ripple band range. Therefore, if no single prominent peak (corresponding to the ripple band) was identified, the event was rejected. Additionally, authentic ripple events were anticipated to display a limited narrowband burst; thus, if the ripple-peak had an excessively wide peak width (indicating broadband spectral changes) or prominent high-frequency activity, the event was considered for rejection (see ^83^).

#### Visual Inspection

Finally, all detected ripples underwent visual inspection, and any events flagged as false-positives during this process were subsequently removed from the dataset.

### Spike sorting

Spike sorting was performed using Kilosort 1.0 (^84^, https://github.com/cortex-lab/KiloSort). The process involved applying a 300-Hz high-pass filter to the raw signals, followed by whitening the data in blocks of 32 channels. Parameters relevant to automated sorting are detailed in the table.

#### Removing putative double-counted spikes

The Kilosort algorithm will occasionally fit a template to the residual left behind after another template has been subtracted from the original data, resulting in double-counted spikes. Such double-counted spikes could artificially inflate inter-spike interval (ISI) violations for a single unit or create erroneous zero-time-lag synchrony between neighboring units. Consequently, spikes with peak times within a 5e-4 second interval and peak waveforms detected on the same channels were systematically removed from the dataset.

#### Removing units with artefactual waveforms

Kilosort1 generates templates of a fixed length (2 ms) that matches the time course of an extracellularly detected spike waveform. However, there are no constraints on template shape, which means that the algorithm often fits templates to voltage fluctuations with characteristics that could not physically result from the current flow associated with an action potential. The units associated with these templates are considered ‘noise’ and are removed on the basis of spread (waveform appears on many channels), and shape (e.g. no peak and trough or sinusoidal waveform) criteria and autocorrelogram function.

#### Manual curation and re-clustering with Phy

Manual curation and re-clustering were performed using Phy (https://github.com/kwikteam/phy). Kilosort-derived clusters were imported into Phy for manual curation. Units that were poorly isolated according to the initial Kilosort results were re-clustered using Klusta with custom-designed plugins (https://github.com/petersenpeter/phyplugins) to obtain well-isolated single units. The quality of these clusters was evaluated based on refractory period violations and Fisher’s linear discriminant metrics. Noise clusters and poorly isolated units were subsequently excluded from the analysis.

### Kilosort Parameters

|  |  |
| --- | --- |
| ops.datatype | 'dat' |
| ops.fs | 30000 |
| ops.Nfilt | 1024 |
| ops.whitening | 'full' |
| ops.nSkipCov | 1 |
| ops.whiteningRange | 32 |
| ops.criterionNoiseChannels | 0.2 |
| ops.Nrank | 3 |
| ops.nfullpasses | 6 |
| ops.maxFR | 20000 |
| ops.fshigh | 300 |
| ops.ntbuff | 64 |
| ops.scaleproc | 200 |
| ops.NT | $32 \times 1024 + \text{ops.ntbuff}$ |
| ops.Th | [4 10 10] |
| ops.lam | [10 30 30] |
| ops.nannealpasses | 4 |
| ops.momentum | $1./[20 \ 400]$ |
| ops.shuffle_clusters | 1 |
| ops.mergeT | 0.1 |
| ops.splitT | 0.1 |
| ops.initialize | 'fromData' |
| ops.spkTh | -4 |
| ops.loc_range | [3 1] |
| ops.long_range | [30 6] |
| ops.maskMaxChannels | 5 |
| ops.crit | 0.65 |
| ops.nFiltMax | 10000 |

Removing units with unstable firing rates: for each neuron, firing rates were estimated in 5-minute epochs throughout the recording duration. During the task, neurons that ceased firing during any trial block were classified as unstable and excluded from the main analyses. Similarly, during sleep, neurons that stopped firing were also deemed unstable and removed from the primary analyses.

### Classification of deep and superficial CA1 pyramidal cells

The full method was previously described in ^50^. We developed a semi-supervised method for classifying cell types based on electrophysiological features of recorded units, specifically single-unit waveforms and interspike interval (ISI) distributions. Before classification, these features were processed using UMAP to reduce the dimensionality of the feature space. The four UMAP-derived features (two from waveforms and two from ISI) constituted the clustering feature space. Spectral clustering, implemented in MATLAB with Mahalanobis distance and k-means, was used to identify clusters.

We identified the cluster with units exhibiting low firing rates, high bursting propensity, and dense localization within the stratum pyramidale as the pyramidal cell group. These physiological features are hallmarks of CA1 pyramidal cells ^85–87^. For superficial and deep analyses, we excluded spatial outliers (units with relative depths outside the 5th–95th percentile range) and cells with mean firing rates >1 Hz or a burst index >3. Since superficial and deep pyramidal cells are classified based on their radial distance within the CA1 pyramidal layer, layer-referenced depth data was required. We used two anterior probes (one per animal) with reliable CA1 sampling across layers.

To estimate linear-array channel locations relative to CA1 layers, we used features of sharp wave ripples (SWRs). The sharp wave component, generated by CA3 Schaffer collateral inputs, produces a current sink in stratum radiatum (SR) and a source in stratum pyramidale (SP), evident as a polarity reversal between these layers. Using current source density (CSD) analysis of ripple-triggered signals, we identified SR and SP regions. The pyramidal layer was further centered using ripple power and the slope of the sharp-wave envelope peak, with the channel at the slope zero-crossing designated as depth 0.

For each session, cells in the “pyramidal” group (meeting firing rate, burst, and depth criteria) were median-split based on depth, creating the superficial group (closer to stratum radiatum) and the deep group (closer to stratum oriens). The spatial distribution of CA1 superficial neurons ranged from −300 to 540 μm, while CA1 deep neurons spanned −630 to −330 μm.

### Decoding of Neural Population with Linear Support Vector Machine (SVM)

The decoding analysis followed the guidelines outlined in the STAR protocols^80^. To decode context information from spiking activity, we estimated the firing rates of each neuron during four task periods for each screen performance: (1) approach to screen, (2) cue, (3) selection, and (4) confirmation. Each period was assigned one of two context labels: Old or New/Recent. Peri-reward periods were excluded from the decoding analysis. Firing rates were normalized (z-scored) to account for variability across neurons, and neurons with unstable firing during task performance were excluded. For each session, a linear SVM was trained to decode context information (Old or New/Recent) using either neural population activity (population decoding) or single-unit activity (single-unit decoding). The “fitcsvm” function in MATLAB was used for classification. For population decoding, population vectors consisting of firing rates across all included neurons were used as features. For single-unit decoding, the firing rate of an individual neuron served as the feature. Class labels were based on the context (Old or New/Recent). The linear SVM was chosen for its optimal generalization performance, even with datasets featuring many variables and limited samples, and its robustness to non-Gaussian distributions^88^. Additionally, the linear SVM provides straightforward interpretability, as the activity of each neuron can be directly associated with its weight in the classification task.

The linear SVM has one hyperparameter, the regularizer, also referred to as the C-parameter. To optimize the performance of the linear SVM, we conducted hyperparameter tuning using a grid search over a range of BoxConstraint values (C = [0.001, 0.01, 0.1, 1, 10, 100, 1000]). The optimal hyperparameters were identified by performing 10-fold cross-validation for each set of hyperparameters. The cross-validation accuracy was calculated as 1 – cvLoss, where cvLoss represents the misclassification rate during cross-validation. We performed hyperparameter tuning for each session independently. Next, we found the range of C-parameter that resulted in best performance for the majority of the sessions. For the main SVM decoding, we used this fixed range for all recording sessions.

For the main SVM decoding, we created a design matrix as described. To refine the hyperparameters within the fixed estimated range, we employed a Bayesian optimization approach, where the BoxConstraint hyperparameter was treated as a continuous variable. The ‘fitcsvm’ function was used with the ‘OptimizeHyperparameters’ option, allowing MATLAB’s Bayesian optimization framework to search the hyperparameter space efficiently. The optimization used the expected improvement acquisition function, and the process was capped at a maximum of 100 objective evaluations. After estimating the optimal hyperparamter within the range, we trained a linear SVM model to decode the context from neural population. We used 10-fold cross-validation to estimate the accuracy of the model.

To assess the statistical significance of the decoding accuracy, we performed permutation test with 5000 permutations. For each permutation, the class labels or firing rates of individual units were shuffled, and the SVM was retrained and tested using the same cross-validation procedure and the hyperparameters of the original model. The p-value was calculated as the proportion of permuted accuracies greater than or equal to the original accuracy. Additionally, bootstrapping (with 10,000 bootstraps) was used to calculate 95% confidence intervals for the permutation accuracies (see Extended Data Fig. 3). To account for the number of units included in the decoder and the imbalance in label ratios, which biased both the main and shuffled SVM accuracies (Extended Data Fig. 3), we designed a shuffling test tailored to each session. Statistical significance was then assessed separately within each session.

To assess the effect of heterogeneity within functional subpopulations where a population is defined by the same sign of the weight, we randomly permuted, without repetition, the vector of firing rates, but only across units with the same sign of the weight. The permutation is done on every trial independently, while labels stay untouched (see Extended Data Fig. 3).

The weight vector (beta coefficients) of the linear SVM, which represents the contribution of each neuron to the classification decision, was extracted from the final model. The weights were normalized by dividing each weight by the L2-norm of the entire weight vector to interpret the relative importance of individual neurons in the classification process and for subpopulation analyses described above.

### Gini coefficient

We computed Gini coefficient which is a commonly used measure of sparsity/selectivity given by:

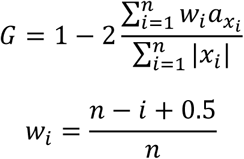

Where *a*_*xi*_ is the absolute values of elements of vector *x* sorted in ascending order, and *n* is the number of elements. We used Typical Sparsity Measures toolbox in MATLAB.

### Assembly pattern identification and activation strength

Cell assemblies were identified during sleep recordings as previously described^31,45,66,89^ (see Extended Data Fig. 4). Significant co-firing patterns were detected using an unsupervised statistical method based on independent component analysis (ICA). The spike trains for each neuron were binned into time windows of 50 ms (corresponding to the 90^th^ percentile of ripple durations) and z-score transformed to eliminate biases due to differences in average firing rates. This creates for each session a cell x binned firing rate matrix (Z). Next, a principal component analysis was applied to the matrix (Z). For this, the correlation matrix of Z was given by 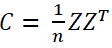 and the eigenvalue decomposition of C was given by:

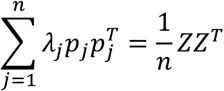

where λ_*j*_ is the j^th^ eigenvalue of C and *p*_*j*_ is its corresponding eigenvector. The Marcenko-Pastur law was used to estimate the number of significant patterns embedded within Z. For a nXB matrix, an eigenvalue exceeding λ_*max*_, defined by 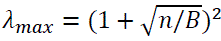, signifies that the pattern given by the corresponding principal component explains more correlation than would be expected if the neurons were independent of each other. The number of eigenvalues exceeding λ_*max*_ was defined as NA and therefore represents the minimum number of distinct significant patterns in the data. The significant principal components were then projected back onto the binned spike data

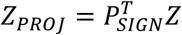

where *P*_*SIGN*_ is the nXNA matrix with the NA principal components as columns.

Independent component analysis (ICA), using the fast ICA algorithm (http://research.ics.aalto.fi/ica/fastica), was then applied to the matrix ZPROJ. That is, an NAXNA unmixing matrix W was found such that the rows of the matrix *Y* = *W*^*T*^*Z*_*PROJ*_ were as independent as possible. The arbitrary signs of the independent component (IC) weights were set so that the highest absolute weight was positive. The unmixing matrix W was then used to derive each cell’s weight within each assembly *V* = *P*_*SIGN*_*W* where the columns of *V* (*i*. *e*., *v*_1_, …, *v*_*NA*_) are the weight vectors of the assembly patterns.

To determine the task activation strength of the assemblies, we tracked each assembly pattern *v_k_*

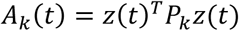

where *Z*(*t*) is a smooth vector function containing for each neuron its z-scored instantaneous firing rate and *P*_*k*_ is the matrix projecting *Z*(*t*) to the activation strength of the assembly pattern *k* at time *t*.

To estimate the reactivation strength of the assemblies detected during task, we used task projection matrices as templates. The spiking matrix of the sleep epoch were binned (50ms), z-scored and projected onto the projection matrices, resulting in reaction strengths that track the time-resolved reactivation of target neuronal ensembles relative to the reference epoch. The diagonal of the projection matrices is set to 0 to make the activation and reactivation strength more robust to the fluctuation of single-neuron activity.

Activation or reactivation vector values greater than 2 times the standard deviation of each epoch independently was used as timestamps of significant expression of each assembly during task or sleep respectively. Re/activation rates were calculated by dividing the number of detected events by the total duration of the epoch of interest.

### Significance test for reactivations

To assess the significance of reactivation events, we performed two separate permutation procedures (Extended Data Fig. 7). No single permutation procedure is able to emulate all possible ways of generating spurious coactivation. Thus, our two approaches were designed to address two different ways of disrupting coactivation with a trade-off between preserving the inter-spike interval (ISI) histograms for each neurons and keeping the simultaneous firing rate modulation intact^1^.

In the first approach, we created surrogate firing rate matrices by shuffling the individual ISIs within the spike train of each neuron. This method preserves inter-spike interval (ISI) histograms for each neuron while compromising the population dynamics and synchronous firing rate fluctuation.

In the second approach, we shuffled the weight of each neuron across the detected templates. In this way, we preserved the ISI histogram of individual neurons and the simultaneous firing of neurons in the firing rate matrix but disrupted the contribution of each neuron to different patterns (i.e. exchanging members across cell assemblies). This is a more conservative approach to testing the reactivation events.

We repeated the permutation process for both approaches 5000 times, and for each permutation, we computed the reactivation vectors based on methods described previously. For each surrogate reactivation vector, we computed the mean of the absolute values and created a surrogate distribution from them. Finally, we compared the mean absolute value of the original reactivation vectors to their corresponding surrogate distributions and estimate the p-value based on a two-sided test.

### Quality of cell assemblies, assembly membership, sparsity, and cosine similarity

Members of candidate cell assemblies were identified using Otsu’s method^33^ to divide the absolute weights into two groups maximizing inter-class variance, and neurons in the group with greater absolute weights were classified as members (Extended Data Fig. 4).

Goodness of separation was quantified using Otsu’s effectiveness metric (OtsuEM), namely the ratio of the inter-class variance to the total variance. Assemblies with small OtsuEM presumably result from limitations of the ICA method in identifying independent components from the PCs^32^. We therefore discarded assemblies with OtsuEM < 0.65 from further analyses.

For each cell assembly weight vector (also referred to as loading or pattern), we computed Gini coefficients. High Gini coefficient (close to 1) indicates high sparsity. In the context of cell assemblies, this means that a small subset of neurons is making a large contribution to the activity of the assembly, while most neurons have very small or no contribution. Thus, the activity is dominated by a few highly active neurons, making the assembly sparse. Low Gini coefficient (closer to 0) indicates low sparsity (or high uniformity). In this case, most neurons contribute similarly to the cell assembly, meaning the activity is more evenly distributed and less sparse. There is a relationship between the Otsu effectiveness metric and Gini coefficients in this context. Because we removed cell assemblies with OtsuEM < 0.65, this affects the lower boundary for the Gini coefficients for all cell assemblies.

To measure the similarity between pair of cell assembly patterns detected in a session, we normalized each pattern to have unit norm and computed cosine similarity using following formula:

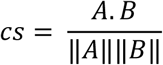

### Assembly coupling

To track interaction between pair of cell assemblies across different conditions, we constructed the assembly coupling matrix which was defined as the Pearson’s correlation coefficient between the re/activation strength vectors.

### Re/activation participation selectivity

For each session, we created a participation probability matrix, N neurons by A assemblies, where each value of the matrix measured how frequently a neuron had participated in re/activation events of a given cell assembly. Participation in each activation event was defined as having fired at least 1 spike. The participation probability of a neuron in a specific cell assembly was calculated as the number of activation events where the neuron fired at least one spike, divided by the total number of detected activation events (Extended Data Fig. 10).

If a cell is highly cell-assembly specific, they would ideally spike with higher proportion in their respective cell assembly and rarely firing during the activation periods of other cell assemblies. To measure the participation selectivity of neurons in cell assembly activation events, we computed Gini coefficient for each row of the participation matrix. A high value of Gini coefficients (closer to 1) show a cell is highly selective. For each cell assembly, member’s exclusivity was defined as the mean of the participation selectivity across identified members of that cell assembly.

This method is related to spike dispersion and tuning to preferred cell assembly in Ref^69^.

### Measuring Bias in Cell Assemblies

Bias in cell assembly activations was assessed using a two-sided binomial test with specified binomial probabilities. For this analysis, we used the MATLAB function ‘myBinomTest’ (Matthew Nelson, 2024. myBinomTest (s,n,p,Sided). Available at: https://www.mathworks.com/matlabcentral/fileexchange/24813-mybinomtest-s-n-p-sided. Retrieved December 2, 2024).

For each cell assembly, significant activation events during task performance were identified as described and categorized based on their occurrence during old and new (Day 1) or recent (Day 2) memory processing. Activations during new trials (s) were defined as successes. To account for differences in the total time spent in trials of varying memory ages, we adjusted the proposed probability of success (p) by weighting it with the fraction of time bins corresponding to new trials.

The ‘myBinomTest’ function computes the probability (pout) of observing the resulting success ratio (s/n) or a more extreme value, given a total of n outcomes and a success probability of p. A cell assembly was classified as old-biased if pout was less than 0.05 and s/n was less than p. Conversely, it was classified as new-biased if pout was less than 0.05 and s/n was greater than p.

### Assembly drift (Kendall’s **τ**)

We used Kendall’s tau correlation coefficient (Kendall’s τ) to estimate the strength and direction of drift in given cell assembly re/activations (similar to Ref ^42^). We defined drift as the monotonic association between re/activation timestamps and re/activation strength. Kendall’s τ is a non-parametric measure that evaluates the correspondence between paired ranks of two variables, providing insight into the ordinal relationship between them. The Kendall’s τ coefficient ranges between –1 and 1, where a value of 1 indicates perfect agreement in ranking, –1 indicates perfect disagreement, and 0 indicates no association. The corresponding p-value was computed to assess the statistical significance of the correlation. We estimated the Kendall’s τ and its significance using the MATLAB function corr, specifying the correlation type as ‘Kendall’. Assemblies with a nonzero slope and p < 0.005 after FDR correction for multiple comparisons across all assemblies were indicated as significant.

### Global– and cell-assembly specific cofiring, connectivity and clustering

For the global measure of cofiring, recordings were segmented into 50 ms time bins, and for each neuron, the number of spikes within each bin was counted and converted to firing rate. Spearman’s rank correlation coefficients were calculated between the firing rate vectors of different neurons, serving as a measure of their co-firing tendencies. We performed this analysis separately for recordings during task, and ripples ripple times during sleep.

Based on the global correlation matrix, we created a weighted adjacency matrix for graph theoretical analysis. The values of the adjacency matrix were equal to the correlation matrix if FDR-corrected p-value of the correlation was < 0.01 and correlation was positive, otherwise weighted adjacency value was set to 0. Thus *a*_*ij*_ and *w*_*ij*_ define whether there existed an edge between nodes *i* and *j*, and what was the strength of the connection.

We computed weighted clustering coefficient given by the following formula:

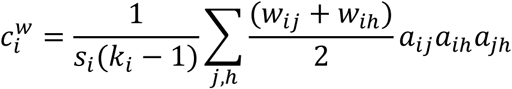

Where 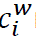 represents the weighted clustering coefficient for node *i*, which quantifies the degree to which node *i* and its neighbors form triangles, considering edge weights. *s*_*i*_: The sum of the weights of edges connected to node *i*, defined as *s*_*i*_ = ∑_*j*_ *w*_*ij*_. This is the total weight of the edges emanating from node *i*. *k*_*i*_: The degree of node *i*, i.e., the number of neighbors of node *i*. *w*_*ij*_, *w*_*i*ℎ_: The weights of the edges connecting node *i* to nodes *j* and ℎ, respectively. These represent the strength of the relationships or interactions between the nodes. *a*_*ij*_: Adjacency matrix elements, where *a*_*ij*_ = 1 if there is an edge between nodes *i* and *j*, and 0 otherwise. The product *a*_*ij*_*a*_*i*ℎ_*a*_*j*ℎ_ ensures that the formula only counts terms where *i*, *j*, and ℎ form a closed triangle.

We defined a metric, cell-assembly-specific cofiring, to capture the degree to which the structure of correlations among neurons contributing to assembly activation changed across time. For each assembly, we extracted the firing rates for all recorded neurons during each assembly activation event. This provides population vectors for each activation event. We then performed Spearman’s rank correlations among the z-scored firing rate of neurons.

For fig 4B, to estimate changes in cell assembly-specific cofiring, we performed the above analysis during early (first 25%) and late (last 25%) activations separately. For each correlation matrix, we computed the mean absolute correlations from the off-diagonal upper part of the correlation matrix. The absolute difference between the early and late mean absolute correlation values provides a measure of overall change in the correlations.

### Superficial-Deep Contribution to New and Old Cell Assemblies

To examine the contribution of superficial and deep pyramidal neurons to the formation of new and old cell assemblies, we used their estimated weights in PCA/ICA space within biased cell assemblies. Since eigenvector (pattern) weights can vary across sessions due to differences in the number of recorded units and the total variance explained by an eigenvector, we developed a robust, session-insensitive measure (Extended Data Fig. 12). First, we normalized the weights within each pattern by dividing each weight by the maximum absolute weight value.

Because individual neurons are not expected to participate in all assemblies within a category, we identified the maximum normalized weight of superficial or deep pyramidal neurons in new or old cell assemblies during a given session, rather than averaging normalized weights. This value is used in Fig. 4F (left).

Typically, multiple superficial or deep pyramidal cells were recorded in a session, resulting in estimates of weights for multiple pyramidal cells per cell assembly. However, as specific pyramidal cells are unlikely to contribute to all cell assemblies, we calculated the maximum weight among all recorded superficial or deep pyramidal neurons to determine the maximum contribution of neurons of a given type to a single cell assembly. This single value per cell assembly is correlated with Kendall’s tau in Fig. 4F (middle).

### Generalized Linear Modeling of Cell Assembly Activity

To quantify how cell assembly activations were modulated by behavioral and contextual variables, we fit generalized linear models (GLMs) to single-trial assembly activation strength values. For each session, we modeled activity from each detected assembly separately. The dependent variable was the mean activation strength of each assembly per trial epoch (e.g. Cue Touch, Decision, Feedback). The following predictors were included in the model: Age context (New, Old sequence type), Screen (ordinal step 1-4 in sequence), Trial Epoch (Approach, Start, Array, Choice), Trial Outcome (Correct, Incorrect), and animal movement metrics (Linear Velocity and Angular Velocity). Age, Screen, Trial Epoch, and Outcome were treated as categorical variables. Interaction terms between Age and both Screen and Trial Epoch were included to capture context-specific modulation of activity.

The model formula was specified as:

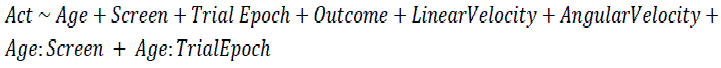

GLMs were fit using MATLAB’s fitglm function with a normal distribution and identity link function.

### Rastermap

We used rastermap^35^ for sorting units in Fig 1D-E for illustration purposes only.

## References

1. Nádasdy, Z., Hirase, H., Czurkó, A., Csicsvari, J. & Buzsáki, G. Replay and time compression of recurring spike sequences in the hippocampus. J. Neurosci. 19, 9497–9507 (1999).

2. Wilson, M. A. & McNaughton, B. L. Reactivation of hippocampal ensemble memories during sleep. Science 265, 676–679 (1994).

3. Gridchyn, I., Schoenenberger, P., O’Neill, J. & Csicsvari, J. Assembly-Specific Disruption of Hippocampal Replay Leads to Selective Memory Deficit. Neuron 106, 291–300.e6 (2020).

4. Lee, A. K. & Wilson, M. A. Memory of sequential experience in the hippocampus during slow wave sleep. Neuron 36, 1183–1194 (2002).

5. Chen, Z. S. & Wilson, M. A. How our understanding of memory replay evolves. J. Neurophysiol. 129, 552–580 (2023).

6. Skaggs, W. E. & McNaughton, B. L. Replay of neuronal firing sequences in rat hippocampus during sleep following spatial experience. Science 271, 1870–1873 (1996).

7. Dave, A. S. & Margoliash, D. Song replay during sleep and computational rules for sensorimotor vocal learning. Science 290, 812–816 (2000).

8. Eliav, T. et al. Fragmented replay of very large environments in the hippocampus of bats. Cell (2025) doi:10.1016/j.cell.2025.05.024.

9. Hoffman, K. L. & McNaughton, B. L. Coordinated reactivation of distributed memory traces in primate neocortex. Science 297, 2070–2073 (2002).

10. Eagleman, S. L. & Dragoi, V. Image sequence reactivation in awake V4 networks. Proc Natl Acad Sci USA 109, 19450–19455 (2012).

11. Umbach, G., Tan, R., Jacobs, J., Pfeiffer, B. E. & Lega, B. Flexibility of functional neuronal assemblies supports human memory. Nat. Commun. 13, 6162 (2022).

12. Vaz, A. P., Wittig, J. H., Inati, S. K. & Zaghloul, K. A. Replay of cortical spiking sequences during human memory retrieval. Science 367, 1131–1134 (2020).

13. Eichenlaub, J.-B. et al. Replay of Learned Neural Firing Sequences during Rest in Human Motor Cortex. Cell Rep. 31, 107581 (2020).

14. Hussin, A. T., Abbaspoor, S. & Hoffman, K. L. Retrosplenial and Hippocampal Synchrony during Retrieval of Old Memories in Macaques. J. Neurosci. 42, 7947–7956 (2022).

15. Sun, W., Advani, M., Spruston, N., Saxe, A. & Fitzgerald, J. E. Organizing memories for generalization in complementary learning systems. Nat. Neurosci. 26, 1438–1448 (2023).

16. Kumaran, D., Hassabis, D. & McClelland, J. L. What learning systems do intelligent agents need? complementary learning systems theory updated. Trends Cogn Sci (Regul Ed) 20, 512–534 (2016).

17. Zaki, Y. et al. Offline ensemble co-reactivation links memories across days. Nature (2024) doi:10.1038/s41586-024-08168-4.

18. Sekeres, M. J., Winocur, G. & Moscovitch, M. The hippocampus and related neocortical structures in memory transformation. Neurosci. Lett. 680, 39–53 (2018).

19. Montefusco-Siegmund, R., Leonard, T. K. & Hoffman, K. L. Hippocampal gamma-band Synchrony and pupillary responses index memory during visual search. Hippocampus 27, 425–434 (2017).

20. Leonard, T. K. & Hoffman, K. L. Sharp-Wave Ripples in Primates Are Enhanced near Remembered Visual Objects. Curr. Biol. 27, 257–262 (2017).

21. Leonard, T. K. et al. Sharp Wave Ripples during Visual Exploration in the Primate Hippocampus. J. Neurosci. 35, 14771–14782 (2015).

22. Karlsson, M. P. & Frank, L. M. Awake replay of remote experiences in the hippocampus. Nat. Neurosci. 12, 913–918 (2009).

23. Carr, M. F., Jadhav, S. P. & Frank, L. M. Hippocampal replay in the awake state: a potential substrate for memory consolidation and retrieval. Nat. Neurosci. 14, 147–153 (2011).

24. Foster, D. J. & Wilson, M. A. Reverse replay of behavioural sequences in hippocampal place cells during the awake state. Nature 440, 680–683 (2006).

25. Tambini, A. & Davachi, L. Awake reactivation of prior experiences consolidates memories and biases cognition. Trends Cogn Sci (Regul Ed) 23, 876–890 (2019).

26. Diekelmann, S. & Born, J. The memory function of sleep. Nat. Rev. Neurosci. 11, 114–126 (2010).

27. Klinzing, J. G., Niethard, N. & Born, J. Mechanisms of systems memory consolidation during sleep. Nat. Neurosci. 22, 1598–1610 (2019).

28. Abbaspoor, S., Rahman, K., Zinke, W. & Hoffman, K. L. Learning of object-in-context sequences in freely-moving macaques. BioRxiv (2023) doi:10.1101/2023.12.11.571113.

29. Chapin, J. K. & Nicolelis, M. A. Principal component analysis of neuronal ensemble activity reveals multidimensional somatosensory representations. J. Neurosci. Methods 94, 121–140 (1999).

30. Peyrache, A., Khamassi, M., Benchenane, K., Wiener, S. I. & Battaglia, F. P. Replay of rule-learning related neural patterns in the prefrontal cortex during sleep. Nat. Neurosci. 12, 919–926 (2009).

31. Lopes-dos-Santos, V., Ribeiro, S. & Tort, A. B. L. Detecting cell assemblies in large neuronal populations. J. Neurosci. Methods 220, 149–166 (2013).

32. Boucly, C. J. et al. Flexible communication between cell assemblies and “reader” neurons. BioRxiv (2022) doi:10.1101/2022.09.06.506754.

33. Otsu, N. A Threshold Selection Method from Gray-Level Histograms. IEEE Trans. Syst. Man Cybern. 9, 62–66 (1979).

34. Jung, B. et al. A comprehensive macaque fMRI pipeline and hierarchical atlas. Neuroimage 235, 117997 (2021).

35. Stringer, C. et al. Rastermap: a discovery method for neural population recordings. Nat. Neurosci. 28, 201–212 (2025).

36. Kudrimoti, H. S., Barnes, C. A. & McNaughton, B. L. Reactivation of hippocampal cell assemblies: effects of behavioral state, experience, and EEG dynamics. J. Neurosci. 19, 4090–4101 (1999).

37. Gorriz, M. H., Takigawa, M. & Bendor, D. The role of experience in prioritizing hippocampal replay. BioRxiv (2023) doi:10.1101/2023.03.28.534589.

38. Dragoi, G. The generative grammar of the brain: a critique of internally generated representations. Nat. Rev. Neurosci. 25, 60–75 (2024).

39. Tingley, D. & Peyrache, A. On the methods for reactivation and replay analysis. Philos. Trans. R. Soc. Lond. B Biol. Sci. 375, 20190231 (2020).

40. Yang, W. et al. Selection of experience for memory by hippocampal sharp wave ripples. Science 383, 1478–1483 (2024).

41. Khatib, D. et al. Active experience, not time, determines within-day representational drift in dorsal CA1. Neuron 111, 2348–2356.e5 (2023).

42. Mau, W., et al. Ensemble remodeling supports memory-updating. BioRxiv (2022) doi:10.1101/2022.06.02.494530.

43. Mau, W., Hasselmo, M. E. & Cai, D. J. The brain in motion: How ensemble fluidity drives memory-updating and flexibility. eLife 9, (2020).

44. Bollmann, L., Baracskay, P., Stella, F. & Csicsvari, J. Sleep stages antagonistically modulate reactivation drift. BioRxiv (2023) doi:10.1101/2023.10.13.562165.

45. van de Ven, G. M., Trouche, S., McNamara, C. G., Allen, K. & Dupret, D. Hippocampal Offline Reactivation Consolidates Recently Formed Cell Assembly Patterns during Sharp Wave-Ripples. Neuron 92, 968–974 (2016).

46. Kossio, Y. F. K., Goedeke, S., Klos, C. & Memmesheimer, R.-M. Drifting assemblies for persistent memory: Neuron transitions and unsupervised compensation. Proc Natl Acad Sci USA 118, (2021).

47. Jenks, K. R., Tsimring, K., Ip, J. P. K., Zepeda, J. C. & Sur, M. Heterosynaptic Plasticity and the Experience-Dependent Refinement of Developing Neuronal Circuits. Front. Neural Circuits 15, 803401 (2021).

48. Fiete, I. R., Senn, W., Wang, C. Z. H. & Hahnloser, R. H. R. Spike-time-dependent plasticity and heterosynaptic competition organize networks to produce long scale-free sequences of neural activity. Neuron 65, 563–576 (2010).

49. Sugden, A. U. et al. Cortical reactivations of recent sensory experiences predict bidirectional network changes during learning. Nat. Neurosci. 23, 981–991 (2020).

50. Abbaspoor, S. & Hoffman, K. L. Circuit dynamics of superficial and deep CA1 pyramidal cells and inhibitory cells in freely moving macaques. Cell Rep. 43, 114519 (2024).

51. Hainmueller, T. & Bartos, M. Parallel emergence of stable and dynamic memory engrams in the hippocampus. Nature 558, 292–296 (2018).

52. Gava, G. P. et al. Organizing the coactivity structure of the hippocampus from robust to flexible memory. Science 385, 1120–1127 (2024).

53. Ólafsdóttir, H. F., Bush, D. & Barry, C. The Role of Hippocampal Replay in Memory and Planning. Curr. Biol. 28, R37–R50 (2018).

54. Foster, D. J. Replay comes of age. Annu. Rev. Neurosci. 40, 581–602 (2017).

55. Girardeau, G. & Lopes-Dos-Santos, V. Brain neural patterns and the memory function of sleep. Science 374, 560–564 (2021).

56. Zhang, J., Ou, J. & Liu, Y. Replay and ripples in humans. Annu. Rev. Neurosci. (2025) doi:10.1146/annurev-neuro-112723-024516.

57. Jang, A. I., Wittig, J. H., Inati, S. K. & Zaghloul, K. A. Human Cortical Neurons in the Anterior Temporal Lobe Reinstate Spiking Activity during Verbal Memory Retrieval. Curr. Biol. 27, 1700–1705.e5 (2017).

58. Griffin, S. et al. Ensemble reactivations during brief rest drive fast learning of sequences. Nature 638, 1034–1042 (2025).

59. Rubin, D. B. et al. Learned Motor Patterns Are Replayed in Human Motor Cortex during Sleep. J. Neurosci. 42, 5007–5020 (2022).

60. O’Neill, J., Senior, T. J., Allen, K., Huxter, J. R. & Csicsvari, J. Reactivation of experience-dependent cell assembly patterns in the hippocampus. Nat. Neurosci. 11, 209–215 (2008).

61. Gupta, A. S., van der Meer, M. A. A., Touretzky, D. S. & Redish, A. D. Hippocampal replay is not a simple function of experience. Neuron 65, 695–705 (2010).

62. Berndt, M., Trusel, M., Roberts, T. F., Pfeiffer, B. E. & Volk, L. J. Bidirectional synaptic changes in deep and superficial hippocampal neurons following in vivo activity. Neuron 111, 2984–2994.e4 (2023).

63. Dupret, D., O’Neill, J. & Csicsvari, J. Dynamic reconfiguration of hippocampal interneuron circuits during spatial learning. Neuron 78, 166–180 (2013).

64. Geiller, T., Fattahi, M., Choi, J.-S. & Royer, S. Place cells are more strongly tied to landmarks in deep than in superficial CA1. Nat. Commun. 8, 14531 (2017).

65. Danielson, N. B. et al. Sublayer-Specific Coding Dynamics during Spatial Navigation and Learning in Hippocampal Area CA1. Neuron 91, 652–665 (2016).

66. Harvey, R. E., Robinson, H. L., Liu, C., Oliva, A. & Fernandez-Ruiz, A. Hippocampo-cortical circuits for selective memory encoding, routing, and replay. Neuron 111, 2076–2090.e9 (2023).

67. Valero, M. & de la Prida, L. M. The hippocampus in depth: a sublayer-specific perspective of entorhinal-hippocampal function. Curr. Opin. Neurobiol. 52, 107– 114 (2018).

68. Davidson, T. J., Kloosterman, F. & Wilson, M. A. Hippocampal replay of extended experience. Neuron 63, 497–507 (2009).

69. Farooq, U., Sibille, J., Liu, K. & Dragoi, G. Strengthened Temporal Coordination within Pre-existing Sequential Cell Assemblies Supports Trajectory Replay. Neuron 103, 719–733.e7 (2019).

70. Cai, D. J. et al. A shared neural ensemble links distinct contextual memories encoded close in time. Nature 534, 115–118 (2016).

71. Lewis, P. A. & Durrant, S. J. Overlapping memory replay during sleep builds cognitive schemata. Trends Cogn Sci (Regul Ed) 15, 343–351 (2011).

72. Tse, D. et al. Schemas and memory consolidation. Science 316, 76–82 (2007).

73. Zhou, Z., Singh, D., Tandoc, M. C. & Schapiro, A. C. Building integrated representations through interleaved learning. J. Exp. Psychol. Gen. 152, 2666– 2684 (2023).

74. Witkowski, S., Schechtman, E. & Paller, K. A. Examining sleep’s role in memory generalization and specificity through the lens of targeted memory reactivation. Curr. Opin. Behav. Sci. 33, 86–91 (2020).

75. Zhao, X., Chen, P.-H., Chen, J. & Sun, H. Manipulated overlapping reactivation of multiple memories promotes explicit gist abstraction. Neurobiol. Learn. Mem. 213, 107953 (2024).

76. Liu, K., Sibille, J. & Dragoi, G. Generative predictive codes by multiplexed hippocampal neuronal tuplets. Neuron 99, 1329–1341.e6 (2018).

77. Liao, Z. et al. Inhibitory plasticity supports replay generalization in the hippocampus. Nat. Neurosci. 27, 1987–1998 (2024).

78. Otero, C., Piza, D. B., Martinez-Trujillo, J. C. & Riera, J. Visual exploration drives Hippocampal SWR rates during 3D spatial navigation in the freely moving marmoset. BioRxiv (2025) doi:10.1101/2025.07.01.662661.

79. Norman, Y. et al. Hippocampal sharp-wave ripples linked to visual episodic recollection in humans. Science 365, (2019).

80. Koren, V. Uncovering structured responses of neural populations recorded from macaque monkeys with linear support vector machines. STAR Protocols 2, 100746 (2021).

81. Talakoub, O. et al. Hippocampal and neocortical oscillations are tuned to behavioral state in freely-behaving macaques. BioRxiv (2019) doi:10.1101/552877.

82. Talakoub, O., Gomez Palacio Schjetnan, A., Valiante, T. A., Popovic, M. R. & Hoffman, K. L. Closed-Loop Interruption of Hippocampal Ripples through Fornix Stimulation in the Non-Human Primate. Brain Stimulat. 9, 911–918 (2016).

83. Chen, Y. Y., Yoshor, D., Sheth, S. A. & Foster, B. L. Stability of ripple events during task engagement in human hippocampus. BioRxiv (2020) doi:10.1101/2020.10.17.342881.

84. Pachitariu, M., Steinmetz, N., Kadir, S., Carandini, M. & Harris, K. D. Kilosort: realtime spike-sorting for extracellular electrophysiology with hundreds of channels. BioRxiv (2016) doi:10.1101/061481.

85. Skaggs, W. E. et al. EEG sharp waves and sparse ensemble unit activity in the macaque hippocampus. J. Neurophysiol. 98, 898–910 (2007).

86. Csicsvari, J., Hirase, H., Czurkó, A., Mamiya, A. & Buzsáki, G. Oscillatory coupling of hippocampal pyramidal cells and interneurons in the behaving Rat. J. Neurosci. 19, 274–287 (1999).

87. Ranck, J. B. Studies on single neurons in dorsal hippocampal formation and septum in unrestrained rats. I. Behavioral correlates and firing repertoires. Exp. Neurol. 41, 461–531 (1973).

88. Belousov, A. I., Verzakov, S. A. & von Frese, J. Applicational aspects of support vector machines. J. Chemom. 16, 482–489 (2002).

89. Peyrache, A., Benchenane, K., Khamassi, M., Wiener, S. I. & Battaglia, F. P. Principal component analysis of ensemble recordings reveals cell assemblies at high temporal resolution. J. Comput. Neurosci. 29, 309–325 (2010).

